# Common Phenomenal and Neural Substrate Geometry in Visual Motion Perception

**DOI:** 10.1101/2025.10.01.679834

**Authors:** Kallum Robinson, Ariel Zeleznikow-Johnston, Jiahao Wu, Yumiko Yoshimura, Naotsugu Tsuchiya

## Abstract

What is a possible physical substrate of the qualitative aspects of consciousness (qualia)? Answering this question is a central goal of consciousness research. Due to their subjective and ineffable nature, finding a quantitative way to characterise qualia from verbal description has thus far proven elusive. To overcome the challenge of expressing subjective experience, recent structural and relational approaches have been proposed from mathematics. Yet, as far as we know, no attempts have been made to evaluate the relationship between a certain structure of qualia and the structure of a candidate underlying physical substrate. Towards this ambitious goal of linking the structures of qualia and physical, we set out to make an empirical first step by focusing on experienced dissimilarity of visual motion in human participants and stimulus-evoked neural population response geometry recorded from mouse primary visual cortex. From human participants (N=171), we obtained dissimilarity ratings of visual motion experiences induced by 48 stimuli, spanning across 8 directions and 6 spatial frequencies. Analysis revealed a human dissimilarity structure that was not well captured by a simple monotonic function of physical motion-direction difference alone nor by a pure orientation-symmetry account. From nine individual mice, we recorded single-neuron activity (n=751) with optical imaging in both awake and lightly anaesthetised conditions (isoflurane 0.6-0.8%). From neuron population responses to a similar set of motion stimuli, we computed a distance matrix that is comparable to our human dissimilarity matrix. Quantitative analyses show structural commonalities between a human dissimilarity structure and mouse neural structure, where a categorical organisation of stimulus direction best explained both. These commonalities were similar in awake and anaesthetised recordings, suggesting that this coarse V1 geometry is relatively insensitive to this type of anaesthesia; future work combining behaviour with causal intervention is required to relate such neural structures to conscious experience. Finally, we list several empirical factors that can be improved to promote our qualia structure approach in the future.

**Graphical Abstract:** Similarity structures derived from a pairwise similarity rating task in humans revealed a mismatch between physical stimuli and subjective experience of similarity of visual motion: participants robustly rated their experience of opposite direction stimuli as similar. In mice, we used 2-photon Ca2+ imaging to record the activity of c.80 neurons per mouse in the primary visual cortex (V1) of nine mice. By computing a correlation matrix that records the similarity of responses to the same pairwise comparisons as in humans across all neurons, we can then represent neuronal ‘distance’. This revealed that mice neurons also exhibited the same dissociation between physical stimuli and responses as in humans at lower spatial frequencies. By comparing these dissimilarity and distance matrices using linear mixed effects modelling, we showed that a categorical model, not direction or orientation, best explains both human and mice data. Ultimately, this study is a methodological proof-of-concept for comparing structures of experience to candidate structures of a possible neural substrate in order to constrain their candidacy.

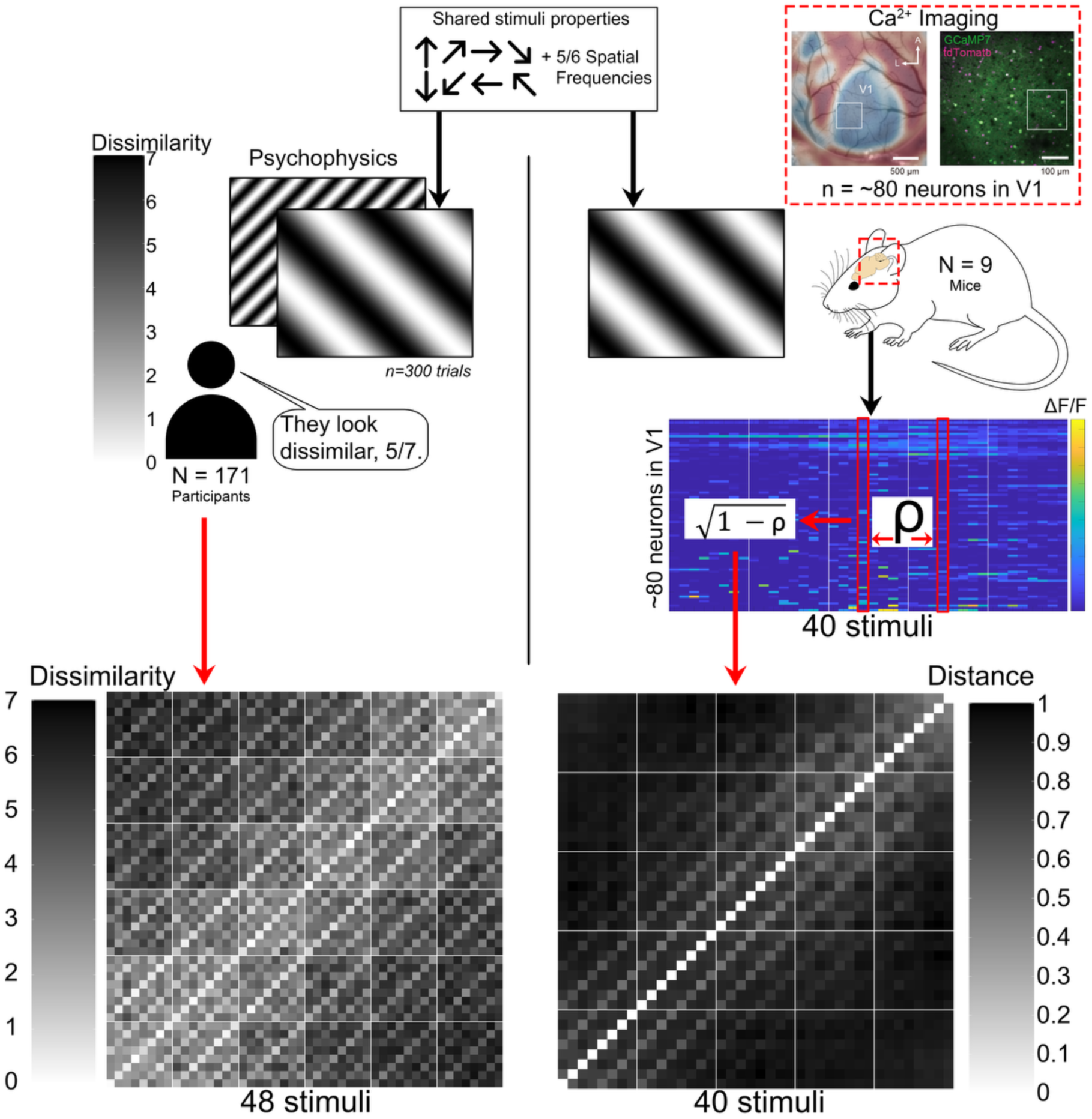

**Highlights:**

- Visual motion feels more similar for opposite than diagonal directions
- Pairwise similarity ratings reveal structure of visual motion experience
- Human motion experience and mouse visual cortex share a common geometry
- Mouse visual cortex codes motion similarly when awake and lightly anaesthetised

## 1. Introduction

Characterising and understanding qualia - the ‘*what it is like*’ to experience something - remains one of the most enduring tasks of modern science (Nagel, 1974; Chalmers, 1995). Throughout this paper, we use *qualia* to refer to the qualitative aspects of consciousness. Any moment of conscious experience, considered qualia in the general sense, is itself composed of multiple *particular qualia*. For example, the experience of standing outside on a summer’s day is itself a quale in the general sense, where the warmth of the sun, the ‘greenness’ of the grass and the sounds of particular insects are examples of qualia in the particular sense. These particular qualia, the specific qualitative aspects of conscious percepts, are the focus of this paper (Balduzzi and Tononi, 2009; Kanai and Tsuchiya, 2012; Lee-Youngzie et al., 2024).

Not only is an understanding of qualia elusive, but also how they are directly associated with brain activity (Lamme, 2006; Koch, 2018). Research seeking an understanding of this association essentially identifies the neural correlates of consciousness (NCC), that is, “the minimal set of neural events jointly sufficient for a conscious state” (Seth and Bayne, 2022). However, in doing so, many studies can lose sight of the nature of qualia itself. One of the reasons is that NCC research tends to categorise qualia as binaries. For example, even if a quale in reality exists on a continuum, studies may ask participants to report it as simply ‘seen’ or ‘unseen’ in order to maximise the inferential power of subsequent contrastive analysis of neural activity (Kob, 2023). While effective, this approach may occlude the intrinsic complexity of qualia and hinder efforts to explain their structure in neural terms (Lepauvre and Melloni, 2021).

To try to overcome this limitation, there is increasing interest within neuroscience, psychology, and computational science in adopting a ‘structural approach’ which aims to go beyond verbal description and instead describe experience in mathematical “language” (Kleiner, 2024; Maier and Tsuchiya, 2026). Mathematical structures such as (generalised) metric spaces offer a flexible framework for capturing both content and dynamics, thus providing more information than contrastive analysis (Lawvere, 1973; Tsuchiya et al., 2022). A metric space consists of a set of points whose pairwise relationships satisfy specific conditions, thus allowing a geometric interpretation of whatever structure these points form in space. Past psychological, neuroscientific, and computational studies have assumed that the target of their respective studies^1^ can be represented as points in some metric space. These studies have investigated the property of the underlying mathematical spaces utilising a general and unifying concept of ‘similarity’ (Shepard, 1987; Kriegeskorte and Kievit, 2013; Roads and Love, 2024). In particular, representational similarity analysis (RSA) (Kriegeskorte and Kievit, 2013) has become popular across different fields of researchers seeking to quantify the relationships between different forms of representations, based on stimulus properties, behaviour, and neural responses (Balkenius and Gärdenfors, 2016; Hebart et al., 2020; Broday-Dvir et al., 2023; Vishne et al., 2023; Kleiner, 2024).

One such structural approach is the ‘Qualia Structure Paradigm’ (Tsuchiya, 2024), which investigates the nature of qualia through relational characterisation – for example, by comparing the similarity between particular qualia. This paradigm is grounded in category-theoretic results (in particular the Yoneda lemma; Tsuchiya and Saigo, 2021; Tsuchiya et al., 2022) showing that a comprehensive description of the relationships among all items in a category is mathematically equivalent to a direct characterisation of the items themselves. Throughout this paper, we use *qualia structure* in the conceptual sense of the Qualia Structure Paradigm, namely as the relational structure taken to characterise a domain of experience. By contrast, in the Methods and Results we refer more directly to the empirically measured *similarity* or *dissimilarity* structure. Humans can quantitatively express their experienced similarity between presented items via pairwise similarity judgements in an experimental setting. A set of stimuli combined with empirically derived similarity ratings forms a structure. By transforming similarity ratings into some distance-like concept, such as dissimilarity, the structure can be treated approximately as a space. In this way, similarity structures derived from similarity judgements over a given class of stimuli can be used to characterise a method-relative proxy of the qualia structure elicited by that class itself. A further clarification of this terminology can be found in the supplementary materials.

An analogous structural approach can also be applied to neural data. Given a set of stimuli, one can compute the pairwise distance between the population response patterns they evoke, yielding a representational dissimilarity matrix (RDM) (Kriegeskorte and Kievit, 2013; Nili et al., 2014). In such analyses, the set of points composing these structures can come from some abstraction of neural activity, as have been measured for example by MEG (Rosenthal et al., 2021), iEEG (Broday-Dvir et al., 2023) or ECoG (Vishne et al., 2023). The central logic is that if an experiential similarity structure and a neural representational structure share common geometry, the neural population may be a candidate substrate for the corresponding domain of experience. Establishing such a structural correspondence between a similarity structure of experience and neural representational geometry is not by itself a demonstration of some substrate instantiating experience. There would remain the strong possibility, given current methods that such correlation could be spurious or statistical. Rather, we treat structural preservation between phenomenal and neural structures as a constraint on selecting further candidate neural substrates of a reported phenomenal experience or domain of experiences. Neural candidates that fail to preserve the relevant structure of experience can be excluded, while those that preserve it can remain plausible candidates (Fink et al., 2021; Maier and Tsuchiya, 2026).

That said, there have been only few efforts to link neural representational geometry to structured domains of experience (Brouwer and Heeger, 2009, 2013; Hirao et al., 2025). If we hope to better understand how subjective experience is supported by a neural substrate, finding links between structures that represent phenomenal experience (derived from human judgements) and structures of neural activity is a promising approach. One limitation of these studies is a reliance on indirect recordings of neural populations in humans. If the critical underlying physical substrate of qualia lies in causal interactions among neurons, then neural activity must be understood in the context of connectivity patterns (Hoel et al., 2013; Haun et al., 2017; Albantakis et al., 2023). However, obtaining direct neural recordings and access to the underlying connectivity structure in humans appears nearly impossible in the foreseeable future, although in vivo recording of very large neural populations per se is becoming a possibility (Quian Quiroga, 2019; Kubska and Kamiński, 2021; Paulk et al., 2022).

In animals, however, it is currently possible to record hundreds of single neurons in vivo and the potential number of simultaneously recorded neurons is rapidly expanding (Hong and Lieber, 2019). Rodents are a common animal model for neuroscience, sharing structural and functional connectivity features with human brains (Xu et al., 2022). Furthermore, it is possible to measure connectivity between neurons in mice both physiologically (Yoshimura et al., 2005) and anatomically (Oh et al., 2014; Whitesell et al., 2021; Sun et al., 2023). Large-scale and long-duration recording also provides enough information to constrain network connectivity within a computational method (Kobayashi et al., 2019). Finally, a recently established behavioural paradigm using pairwise-similarity in mice is very promising in its demonstration of a metric space that reflects mice’s perceptual distance between odours (Nakayama et al., 2022). However, this behavioural approach in mice is time-consuming and only primarily used to resolve identical and non-identical stimulus pairs.

Therefore, we presently have no perfect model system where we can simultaneously obtain direct reports about qualia (as is possible in humans) and obtain relevant connectivity and activity states of neurons at single-cell resolution (as is presently possible in many animal models). The pathway through animal models is likely to make steady progress, although tracing the connectivity of neurons and training animals to report percepts are very time- and resource-consuming. A pragmatic way forward is to combine these complementary strengths. This builds on a long tradition of relating psychophysics to neural measurements in animals, notably in motion research on non-human primates using lesions, single-unit recordings and later causal manipulations (Newsome and Pare, 1988; Newsome et al., 1989; Parker and Newsome, 1998), as well as on more recent mouse psychophysics platforms combining imaging with perturbation (Burgess et al., 2017; Gale et al., 2024). Here then, we aim to extend these lines of work by estimating a large human dissimilarity structure from explicit pairwise experiential judgements over drifting grating motion stimuli, and comparing it directly with a stimulus-evoked neural population dissimilarity structure derived from matched mouse recordings. Human psychophysics provides access to reported experiential similarity, while mouse recordings provide access to large neural populations and, in principle, their underlying connectivity. In this sense, the present study serves as a preliminary first step toward characterising how estimated qualia structures from humans may relate to neural population structures. This relates to similar representational-space approaches comparing behavioural and neuronal similarity structure, such as work on macaque inferotemporal neurons (Op De Beeck et al., 2001), later representational dissimilarity frameworks (Kriegeskorte, 2009) and more recent within-species work in behaving macaques has linked neuronal activity to perceptual judgements in structured stimulus domains, although not via large pairwise similarity matrices of the type used here (Clark and Bradley, 2022; Ziemba et al., 2024).

In this paper, we focus on the phenomenology induced by drifting sinusoidal gratings. For this type of stimulus, motion direction is intrinsically coupled to the orientation of the grating. Accordingly, throughout this paper we use ‘visual motion qualia’ to refer to the experienced similarity structure elicited by these gratings where direction and orientation are coupled together. Visual motion is a very well empirically studied aspect of vision, stemming from studies utilising classical psychophysics paradigms dominated by discrimination thresholds (Levinson and Sekuler, 1975; Nishida, 2011). There are also efforts to dissociate aspects of external motion stimuli from conscious experience by utilising ambiguous motion (Wallach, 1935; Kane et al., 2011) and illusory motion (Braddick et al., 2002; Kitaoka and Ashida, 2003; Takemura et al., 2011). However, as far as we are aware, the quality of visual motion stimuli has not yet been characterised by similarity rating tasks, especially at the scale we present here. We theorise that this is due to an assumption that visual motion perception is necessarily ‘veridical’ and unlikely to contain an interpretable element like that, say, of colour perception.^2^ Perhaps one reason is that much of the visual motion literature has focused on discrimination, estimation and veridical reporting of motion direction, rather than the qualitative similarity between motion experiences <u>(Levinson and Sekuler, 1975; Nishida, 2011)</u>.

Therefore, in meeting the need for an understanding of the structure of visual motion qualia in humans, this study collected pairwise similarity ratings of visual motion stimuli of 8 different directions across 6 spatial frequencies from 171 participants in addition to a series of control studies. We directly compared this similarity structure to a distance structure estimated from neural population activity patterns recorded via two-photon calcium imaging of the primary visual cortex (V1) from 9 mice with an original purpose not directly relevant to this study (Wu et al., 2026). Around 80 responsive neurons were analysed from each mouse while they were awake and then under anaesthesia with their eyes held open. Some mice were also recorded while awake again after recovery from anaesthesia, confirming the identity of individual neurons over a long period of recording. There is evidence that V1 contributes importantly to visual processing and visually guided behaviour in awake mice (Muir et al., 2015; Palagina et al., 2017; Marques et al., 2018; Goldbach et al., 2021). Therefore, under the light anaesthesia used here, neural responses in V1 are thought to remain largely intact whereas the status of conscious visual experience is uncertain. We therefore expected that if the anaesthetic manipulation disrupted neural structures more closely related to conscious visual processing, the neural activity structure obtained under anaesthesia would show reduced commonality with the human dissimilarity structure relative to the awake condition.

## 2. Design and Methods

### 2.1 Psychophysics Design

#### 2.1.1 Participants

A total of 200 participants were recruited for psychophysics experiments through the online platform ‘Prolific’ (https://www.prolific.com). Prolific recruits participants from a global pool and is a respected source of high-quality data (Douglas et al., 2023). Participants were screened for English fluency in order to understand the instructions and that they ran the experiment on a desktop computer. Participants then provided some basic demographics. Each participant was paid at a rate of 9£/hr with a median time to complete the experiment of 25 minutes. Participants enrolled in the main experiment or one of the three smaller control experiments described later. None participated in more than one experiment, including any pilot version of the experiment, to reduce any effect of prior exposure to the similar tasks.

#### 2.1.2 Online Experiment

All psychophysics experiments were developed using a custom PsychoPy script (v. 2023.2.3) (Peirce et al., 2019) run in PsychoJS and hosted online via the ‘Pavlovia’ (https://pavlovia.org) platform. Participants recruited via Prolific were directed towards Pavlovia to run the experiment via an internet browser in full-screen. As this task is administered online, we cannot fully control the conditions under which participants performed the experiment. The reliability of PsyhoPy in conjunction with online platforms have been documented (Leadbeater et al., 2023; Flanagan et al., 2025) and we have successfully used them in similarity judgements to assure equivalent results between in-lab and online participants directly (Moriguchi et al., 2025) (Also see (Qianchen et al., 2022)). In this experiment, we did not make particular efforts to precisely control viewing distance, physical screen size, browser-scaling or ambient room conditions, as they are not very relevant to our paradigm (see (Zeleznikow-Johnston et al., 2023) on how to control these aspects for colour similarity experiments). The experiments in this paper focus on the relational structure of similarity judgements, in which the quality of the data is best assured through the characterisation of the response distributions within a given participant. In this paper, we used double-pass correlation (i.e., internal consistency between the first and the second similarity judgements to the same stimulus pair) and mean-standard deviations of all responses (i.e., removing participants who did not distribute similarity responses as requested in the instruction). In similarity experiments as in this paper, these screening procedures ensure the quality of the data, making it as reliable as the directly compared in-lab collected data (Moriguchi et al., 2025).

#### 2.1.3 Stimuli

Stimuli were generated through PsychoPy’s ‘gratingstim’ function. Sinusoidal gratings were presented full-screen at a temporal frequency of 2Hz. Assuming that the average distance between participants’ eyes to the screen is 70 cm with a screen width of 35cm (corresponding to a 16” diagonal screen), 1 degree visual angle corresponds to 1.2cm. Thus, the spatial frequencies shown for the online psychophysics experiment are *∼*0.04cpd, 0.07cpd, 0.14cpd, 0.29cpd, 0.58cpd and 1.16cpd. The direction of the grating’s movement varied by 8 increments of 45°, from 0° to 315°.

#### 2.1.4 Main experiment

Once participants opened the experiment, they were notified that the estimated duration of the experiment would be roughly 30 minutes and that they were free to quit at any time by pressing the escape key. Following this, participants saw an informed consent page approved by the Monash University Human Research Ethics Committee (Project ID: 17674 and 41190).

Participants were then provided with written instructions on how to complete the experiment along with a short video displaying what to expect and how to select an answer. Participants were instructed to respond how similar the two gratings were using an integer from −4 to +4 (excluding 0) where −4 represents “very different” and +4 represents “very similar”. These similarity judgements were collected using the eight-option radial response interface in Fig.1C. This type of response interface reduces known response biases induced by other configurations (see Supplementary Methods). At the beginning of the experiment, participants were shown a set of 16 motion stimuli (within a square of 200px in length and width) so that they can form a rough idea of the range and variation of stimulus characteristics they should expect. These 16 stimuli covered all 8 possible directions of motion and all possible 6 spatial frequencies in a single random combination shown to all participants. A full recording of these instructions can be found at <u>(Robinson, 2026).</u>

**Fig. 1.**
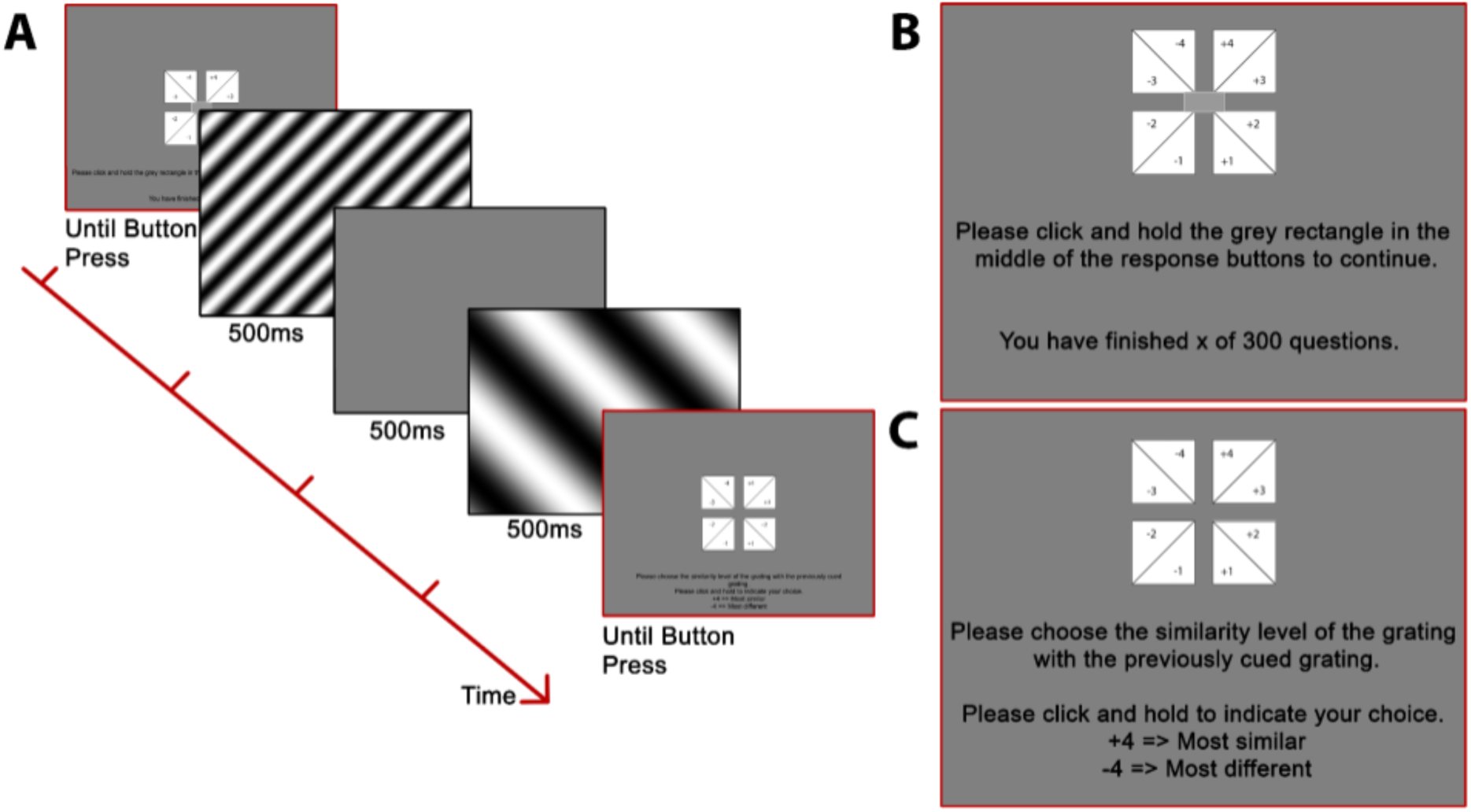
Experimental procedure. **A.** The two screens highlighted in red (first and last) are expanded in **B** and **C** to enhance readability of instructions in this figure. **B.** Participants began a trial by clicking a small rectangle in the centre of the screen. The first stimulus was presented for 500ms followed by a 500ms inter-stimulus interval then by the presentation of the second stimulus for 500ms. **C.** Participants then indicated the similarity between two stimuli from the range of −4 (most different) to +4 (most similar) omitting 0, without time restriction.

Participants then completed 7 practice trials, with 2 stimuli presented in each trial. These 14 stimuli were the same across all participants, in terms of the order of presentation, spatial frequency, and direction characteristics. These practice trials included feedback on their responses. Upon a selection of −4 or −3, the participants saw “you found the experience of the two moving gratings very different” on their display for 4 seconds. Conversely, upon a selection of +4 or +3, they saw “you found the experience of the two moving gratings very similar.” If participants selected +1 or +2, they were told “you found the experience of the two moving gratings similar.” Conversely, if they selected −1 or −2, they were told “you found the experience of the two moving gratings different.” Following completion of these practice trials, main trials proceeded according to Fig. 1A.

#### 2.1.5 Trial Distribution

Given 8 directions and 6 spatial frequencies, this study tested 48 different combinations of stimuli parameters, resulting in 2304 pairwise comparisons to be made. Once participants had enrolled in the experiment in Prolific, they were assigned a subset of 150 stimuli pair combinations, which were presented to them as a sequence in a random order. Due to a software error, the sampling of stimuli pairs was not fully uniformly distributed across participants (min = 12, median = 21, max = 38, see Supplementary Fig S1. & Supplementary Methods).

The order of stimulus combination from 1st to 150th trial (the first pass) was duplicated exactly for 151st to 300th (the second pass). Within a given trial in the second pass, the sequence of the pair of stimuli was reversed with respect to that of the first pass. In total, each participant had 300 trials to complete where the median time for the whole experiment was 25 minutes. Participants were not informed of the design of the double pass paradigm, and there was no explicit break between the first and the second pass (Supplementary Fig. S1).

#### 2.1.6 Exclusion

We excluded participant data by three pre-set criteria before conducting the analysis. First, we excluded participants who failed to complete the entire experiment. 7 out of 200 failed to complete, leaving 193 participants. The second criterion is the Spearman correlation ρ between two sets of similarity ratings given to the first and the second pass as described above. We excluded participants whose correlation between the first and second pass (double pass correlation ρ), was lower than 0.1603. This value was calculated according to the critical ρ value for n=150 (the number of pairwise trials) with an alpha of 0.05. Low double pass correlation is indicative of neglect or inattention to the experiment (Awwad Shiekh Hasan et al., 2012). 13 participants scored below this cutoff and were excluded from further analysis, leaving 180 (93%) (Fig. 2A). Finally, participant responses were analysed for outliers (2 standard deviations) in variance of similarity judgements. Outliers in variance may still have high double-pass scores and need to be considered individually (Fig. 2B). High variance of responses indicates participants provided answers only at the extremes of the spectrum (−4 or +4) and were less likely to be giving representative responses. 9 participants’ similarity responses were outliers and were excluded, leaving 171 participants for all further analysis. Outliers in mean similarity (n=10) could still indicate a particular phenomenal experience and were left in the pool.

**Fig. 2.**
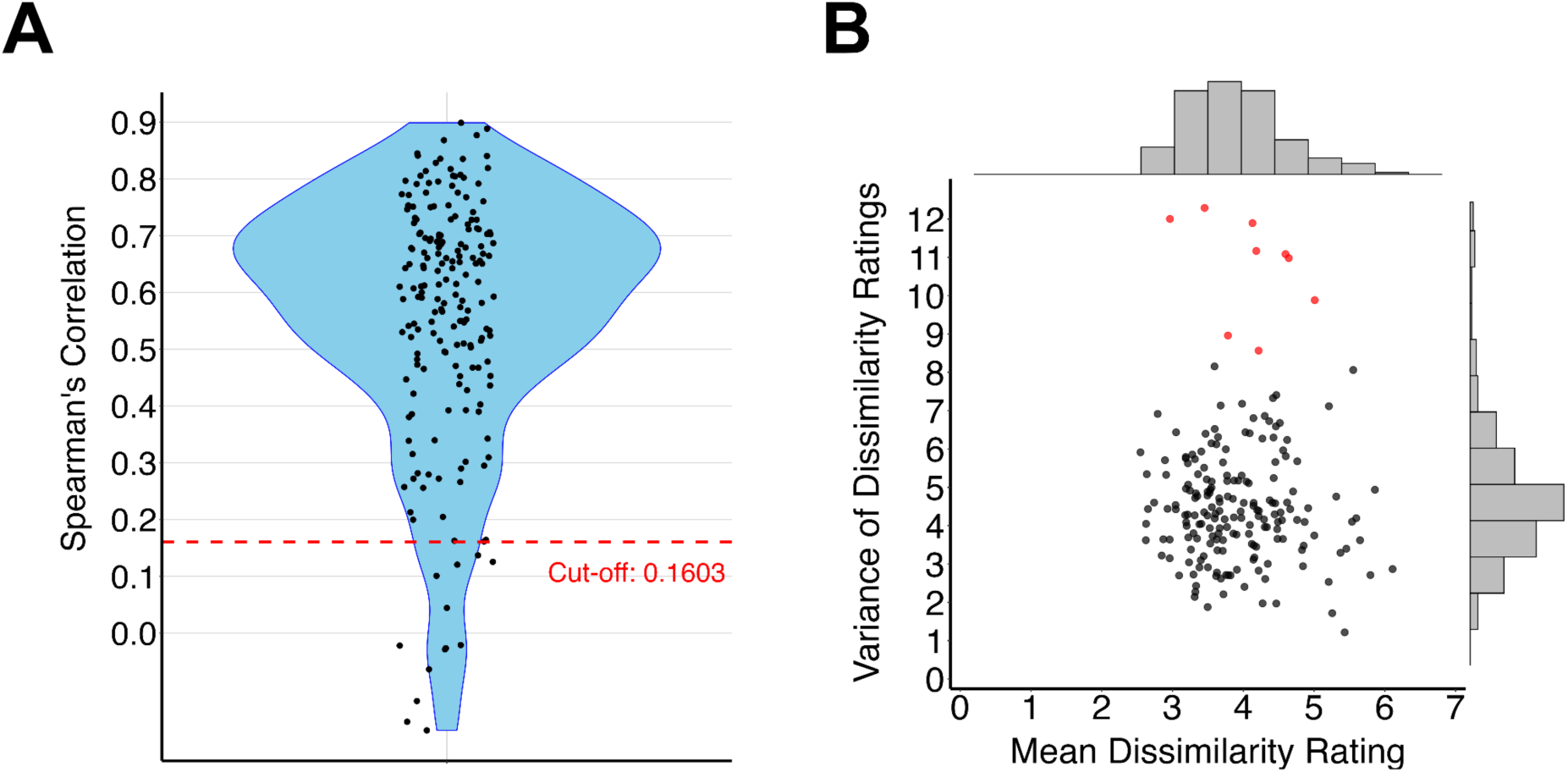
Participant exclusion criteria. **A)** Each point represents an individual participant’s (N=193) Spearman correlation coefficient between the two trial blocks each consisting of 150 trials. Red dashed line denotes cut-off calculated from critical ρ value for n=150 (0.1603). **B)** In order to individually evaluate participant performance, we calculated average dissimilarity rating across all trials and variance of ratings across all trials for individual participants. Points represent all individual participants (n=193). Red points denote outliers (> 2 s.d.) in variance who were excluded from further analysis (n=9). Box plots describe distribution of participants in each axis.

**Fig. 3.**
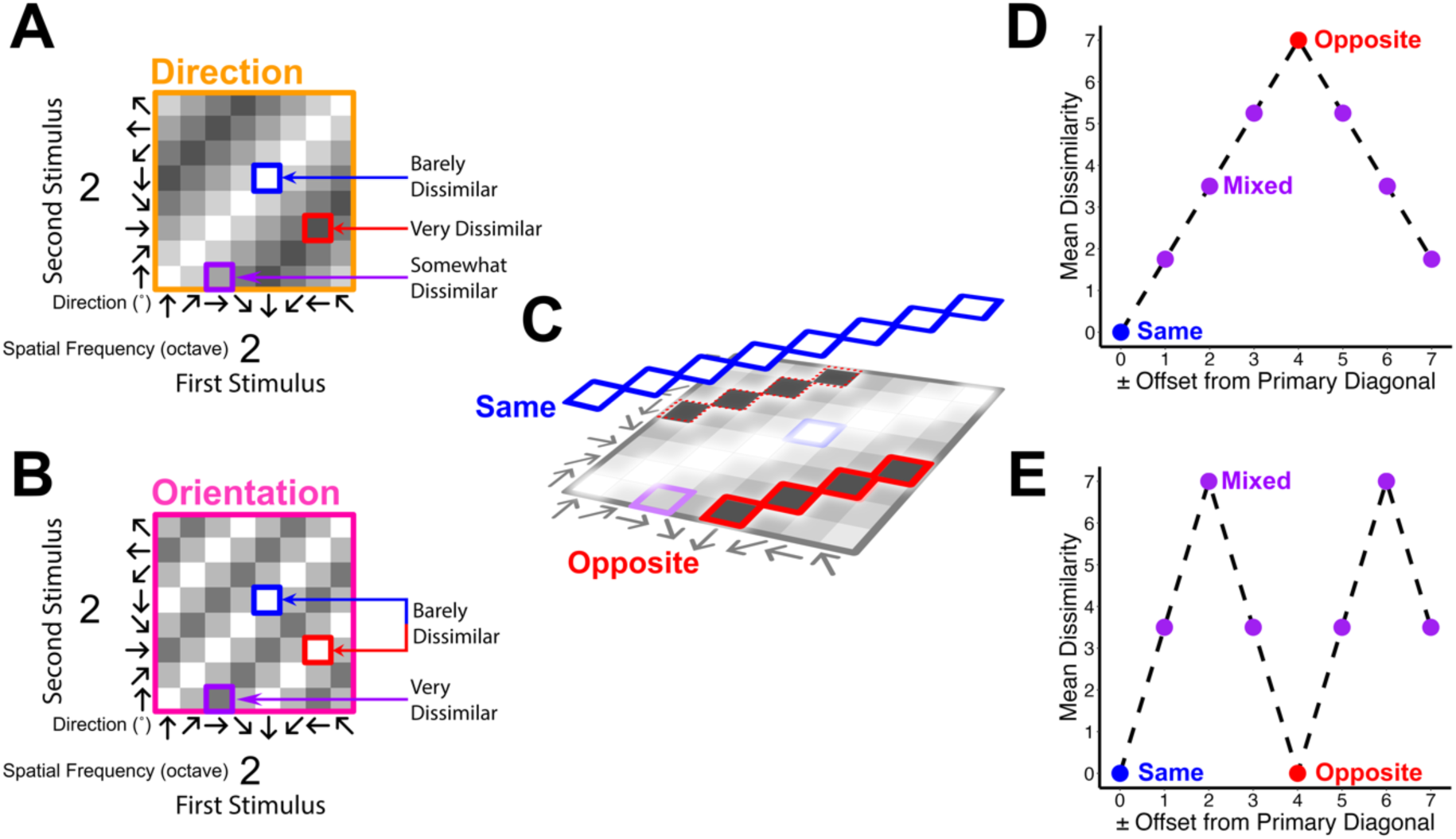
Idealised dissimilarity matrices and offset plots for Direction and Orientation representational models. These models are shown as schematic references to guide later interpretation of the empirical results, they are not experimental data. **A & B** are idealised dissimilarity matrices showing what Direction (orange) and Orientation (pink) representational models would predict within a single spatial frequency octave, respectively (see section 2.3.4). Blue boxes and arrows mark identical-direction stimuli, red marks opposite-direction stimuli and purple marks to all other mixed cases. **C** shows how these matrices are used to make the diagonal offset plots in D and E, and throughout the paper. In the case of identical stimuli (blue diagonal), we take an average along this diagonal with an offset of 0. In the case of opposite direction stimuli (red diagonal) we take an average along this diagonal with an offset of +4 (solid line), as well as −4 (dotted line) and group them together. The same is done for mixed diagonals (not shown). **D & E** shows how these diagonal averages are represented on a diagonal offset plot as points for both direction and orientation representational models, respectively. In an ideal direction model (D), opposite directions are maximally dissimilar from the same direction. In an ideal orientation model (E), same and opposite direction pairs share the same carrier orientation and are thus similarly identical.

#### 2.1.7 Control experiments

We conducted three control experiments to test the generality of the opposite-direction similarity effect observed in the main experiment. All three control experiments shared the following design: motion directions were sampled in 18° increments from 0° to 342° (20 directions), spatial frequency was fixed at 0.16 cpd, and stimuli were presented within a circular aperture (200 px diameter, ∼6° visual angle) appearing in one of four quadrants (∼5° from fixation, randomised across trials and participants). Stimulus duration was 100 ms with a 250 ms interstimulus interval. Each participant completed all 210 pairwise direction combinations twice (420 trials total), with presentation order reversed in the second block for double-pass correlation. Twenty participants were recruited for each experiment via Prolific as per 2.1.1, with a double-pass correlation threshold of 0.139 (critical r for 200 observations, α = 0.05).

The three experiments differed only in temporal frequency: 0.3 Hz, 2.4 Hz, and 18 Hz. These values were chosen to span a range from very slow motion (0.3 Hz, where the stimulus appears near-static) to very fast motion (18 Hz, where only the grating orientation is visible). After exclusions for incomplete data and low double-pass correlation, the final samples were N = 16 (0.3 Hz), N = 17 (2.4 Hz), and N = 17 (18 Hz). Instructions and practice trials followed the same procedure as the main experiment (section 2.1.4).

To directly assess the contribution of carrier orientation to the observed dissimilarity structure, we conducted an additional control experiment using static oriented gratings. The experiment was identical to the main experiment (section 2.1.4) in all respects — stimulus parameters (8 orientations × 6 spatial frequencies = 48 conditions), trial distribution, double-pass design, exclusion criteria, and instructions — except that the gratings did not move. Participants (N = 81), recruited via Prolific as per section 2.1.1) rated the pairwise similarity of static sinusoidal gratings varying in orientation (0° to 315° in 45° steps) and spatial frequency (0.04 to 1.16 cpd). In this stimulus set, orientations separated by 180° are physically identical (e.g., a 0° and 180° static grating are the same horizontal grating), whereas in the main experiment with drifting gratings, 0° and 180° correspond to opposite directions of motion along the same axis. The resulting dissimilarity matrix was constructed as per section 2.3.1. To isolate the effect of motion on the similarity structure, we computed a cell-wise difference matrix by subtracting the main experiment’s drifting-grating dissimilarity matrix from the static-grating dissimilarity matrix (i.e., static − moving). Positive values indicate pairs rated as more dissimilar for static than moving gratings; negative values indicate pairs rated as more similar for static gratings. A summary of all stimulus and design parameters across the main experiment, control experiments, and mouse study is provided in Table 1.

**Table 1.** Summary of stimulus and design parameters across all experiments. Stimulus and design parameters for the main human experiment, four human control experiments, and the mouse study. Spatial frequencies for the main experiment are approximate values assuming a 70 cm viewing distance (see section 2.1.3). The mouse study used 5 spatial frequencies in one-octave steps from 0.02 to 0.32 cpd. N/A = not applicable. ISI = interstimulus interval. cpd = cycles per degree.

| Parameter | Main Experiment | Control 1 | Control 2 | Control 3 | Orientation control | Mouse study |
| --- | --- | --- | --- | --- | --- | --- |
| **N (analysed)** | 171 | 16 | 17 | 17 | 81 | 9 |
| **Directions** | 8 (45° steps) | 20 (18° steps) | 20 (18° steps) | 20 (18° steps) | 8 (45° steps) | 8 (45° steps) |
| **Spatial frequencies** | 6 (0.04–1.16 cpd) | 1 (0.16 cpd) | 1 (0.16 cpd) | 1 (0.16 cpd) | 6 (0.04–1.16 cpd) | 5 (0.02–0.32 cpd) |
| **Temporal frequency** | 2 Hz | 0.3 Hz | 2.4 Hz | 18 Hz | 0 Hz | 2 Hz |
| **Stimulus type** | Full-screen grating | Circular aperture (~6° diam.) | Circular aperture (~6° diam.) | Circular aperture (~6° diam.) | Full-screen grating | Full-screen grating |
| **Stimulus location** | Central | One of 4 quadrants (~5° eccentric) | One of 4 quadrants (~5° eccentric) | One of 4 quadrants (~5° eccentric) | Central | Contralateral full-field |
| **Display extent** | ~28° azimuth (est.) | ~6° diameter | ~6° diameter | ~6° diameter | ~28° azimuth (est.) | ~100° azimuth |
| **Stimulus duration** | 500 ms | 100 ms | 100 ms | 100 ms | 500 ms | 2000 ms |
| **ISI** | 500 ms | 250 ms | 250 ms | 250 ms | 500 ms | 4000 ms |
| **Unique stimulus pairs** | 2304 (distributed) | 210 (within-subject) | 210 (within-subject) | 210 (within-subject) | 2304 (distributed) | N/A (passive viewing) |
| **Trials per participant/mouse** | 300 | 420 | 420 | 420 | 300 | 10 repeats × 40 conditions |
| **Task** | Similarity rating (–4 to +4) | Similarity rating (–4 to +4) | Similarity rating (–4 to +4) | Similarity rating (–4 to +4) | Similarity rating (–4 to +4) | Passive viewing |

### 2.2 Mice Study Design and Data Acquisition

Authors JW and YY conducted this portion of the study prior to, and independent of, the human psychophysics study (Wu et al., 2026). All animal experiments were approved by the Experimental Animal Committee of the National Institute for Physiological Sciences, Japan. The main motivation and aim of the mice experiment is irrelevant to the current paper. The methodological details are reported in (Wu et al., 2026) and here we provide only necessary details to understand this paper.

To label Vgat-positive interneurons, Vgat-IRES-Cre mice (The Jackson Laboratory, stock #16962) were crossed with Ai14 tdTomato reporter mice (The Jackson Laboratory, stock #007908). Mice of either sex aged 2–5 months were used. To express calcium indicator jGCaMP7, AAV1-Syn-jGCaMP7b-WPRE (Addgene) was injected into the primary visual cortex of the mice anaesthetised by intraperitoneal injection of medetomidine (0.75 mg/kg; Domitor, Zenoaq), midazolam (4.0 mg/kg; Midazolam sand, Sandoz), and butorphanol (5.0 mg/kg; Vetorphale, Meiji Seika). A cranial window was implanted 10–12 days after the virus injection under the same anaesthesia as above. These surgical anaesthesia were unrelated to the experimental anaesthetic condition (see below). All animals were recovered in homecage for three days before undergoing neural recording.

Mice heads were fixed to a custom stereotaxic instrument for two-photon calcium imaging. GCaMP fluorescence was acquired using a two-photon microscope (A1R MP, Nikon) equipped with a Mai Tai DeepSee laser. GCaMP and tdTomato were excited at 950nm using 500/50nm and 663/75nm emission fibres, respectively. Imaging depth was set at 190-300μm (presumed to be layer 2/3).

Critically, each of nine mice was shown a series of full-screen sine-wave drifting gratings in a similar way to the human psychophysics. In each trial, the grating was presented at 2Hz temporal frequency, then randomised by some combination of 8 directions (0° to 315° in 45° steps) and 5 spatial frequencies (0.02cpd to 0.32cpd in one-octave steps). Notable differences between the mice and the human experiments are as follows. First, the stimulus duration was longer for mice (2 seconds) than humans (500ms). Second, mice did not make any perceptual judgments and viewed the stimuli passively in the anaesthetised and awake state, with a fixed interstimulus interval (4s). For humans, the interstimulus interval was 500ms and the initiation of the next trial was self-paced (Fig. 1). Humans were only tested in the awake state. Third, the display size extended over 100 degrees in azimuth of visual field for the mice but covered an estimated average of 28 degrees in azimuth of visual field for humans.

For mice, GCaMP fluorescence was recorded by cellular-level two-photon calcium imaging across 10 trials. Upon preliminary responsiveness check (See below, Wu et al., 2026), we analysed approximately 80 neurons from each mouse’s V1 by computing the mean fluorescence level during the 2s stimulus window. Mice were first imaged in the awake condition and then under light isoflurane anaesthesia. Isoflurane levels were >1.2% for induction and 0.6-0.8% in air for maintenance while their eyes were held open. 0.6-0.8% isoflurane in air was regarded as deep enough anaesthesia to guarantee loss of behavioural responsiveness in mice. An even deeper level of isoflurane would reduce the amplitude of visual stimulus-evoked responses and cause slow rolling eye movements (Ikeda and Wright, 1974; Schummers et al., 2008).

All neurons included in the analyses were those that have clear responses in a time locked manner with respect to visual stimuli for both wake and anaesthetised conditions (Wu et al., 2026). Across 9 mice, we identified 751 neurons, with 626 excitatory and 125 inhibitory neurons in total. Orientation and direction tuning properties of these neurons are summarised in Supplementary Methods and are explained in further detail in the Wu and colleagues’ paper. For all analysis, we made no distinction between excitatory and inhibitory neurons. We defined the visual response magnitude in each trial as the average of ΔF/F from the onset to the offset of the stimulus duration (over 2 seconds), normalised to the baseline averaged from the 1.3s prior to stimulus presentation within the 4 seconds of interstimulus interval.

### 2.3 Analysis

#### 2.3.1 Psychophysics Dissimilarity Matrices

We performed all analyses using R Studio (v.2024.12.1+563). To reduce the confusion among subjects (possible flips of response mapping between similar and dissimilar, which we observed for many online participants (Zeleznikow-Johnston et al., 2023), we explicitly used negative ratings (−4 to −1) for “different” and positive ratings (+1 to +4) for “similar”. Before conducting analysis, we first converted the similarity rating scores of −4, −3, −2, −1, +1, +2, +3, +4 into dissimilarities of 7, 6, 5, 4, 3, 2, 1, 0 respectively in order interpret these ratings as distance-like entity with a unit interval. Due to the distributed study paradigm, no single participant saw all 48 × 48 combinations. Therefore, dissimilarity scores were compiled in a matrix as an average of all participants’ similarity ratings. Mean dissimilarity was calculated for each stimulus combination by aggregating all participants by stimulus condition. Each column and row represent a unique stimulus parameter combination. Reported values are given as dissimilarity ±1 standard deviation.

#### 2.3.2 Neural Similarity Response and Distance Matrices

After averaging across time and trials, we obtained ΔF/F data as a 3D matrix, with each dimension being neuron ID, direction, and spatial frequency. To define the similarity of neural response patterns for each combination of motion direction and spatial frequency, we computed Spearman’s correlation between a vector of the averaged response over N neurons. For example, for a particular session with 80 neuron recordings, we had a 80×1 dimensional vector per each stimulus combination. To construct a neural response similarity matrix, we used 1-r_ij_, with r_ij_ being the Spearman correlation coefficient between the vectors from the stimulus combination of i and j. Next, we converted 1-r_ij_ to √1 − *r_ij_* which is a metric (Solo, 2019) and then set the diagonal to 0 (Supplementary Fig. S2). Bootstrap analysis with 10,000 iterations was performed to estimate variability and calculate standard deviation of distance. Reported values are given as distance ± 1 standard deviation. The resulting distance values off the main diagonal, prior to any rescaling, spanned a range of approximately 0.28 to 1.25 across all mice in the awake condition against a theoretical maximum of √2. This indicates a well-differentiated spread of neural response pattern dissimilarities across stimulus conditions. To enable visual comparison between mice and human matrices in figures,we applied a min-max rescaling of distance values to an interval of [0, 1]. This monotonic linear transformation preserves the rank order and relative spacing of all pairwise distances. These rescaled values were not used for any analysis (see section 2.3.5).

#### 2.3.3 Multidimensional Scaling (MDS)

MDS takes a dissimilarity matrix and returns a set of points in space, such that the distance between points represents the similarity between individual cases of a data set. MDS is a useful tool for visualising high-dimensional data by reducing dimensions to a level that can be visualised. To perform the multidimensional scaling visualisation (Borg and Groenen, 2005), we first symmetrised the dissimilarity matrix by adding the transpose of the dissimilarity matrix to itself and dividing by 2. This symmetrisation removes order-specific asymmetries in sequential judgements so that subsequent analyses reflect the shared pairwise structure rather than A→B versus B→A presentation effects. We then also replaced the diagonal entry with 0.

As human data contains violations of the standard Euclidean metric, all human MDS was performed using Kruskal’s non-metric MDS via the ‘smacof’ function in R (Mair et al., 2022). Kruskal’s non-metric MDS focuses on preserving the rank order of dissimilarities and is robust to non-linear scaling, which is ideal when dissimilarity ratings are ordinal, as in our case.

For mice data, we used the ‘cmdscale’ function in R, which follows the analysis of (Mardia, 1978), returning the best fitting k-dimensional metric representation in Euclidean space.

#### 2.3.4 Idealised model matrices and diagonal offset summaries

To aid interpretation of the empirical dissimilarity matrices, we constructed two idealised models of dissimilarity between two stimuli in terms of how they differ either by direction or orientation.

For the direction model, predicted dissimilarity increased linearly with the circular angular difference between the two stimulus directions. Angular differences were first wrapped onto the range 0° - 180°, such that 0° indicated identical directions and 180° opposite directions. These values were then linearly scaled onto the dissimilarity range 0-7. Such a model captures the simplest monotonic expectation if human judgements and neural responses varied directly with physical motion-direction difference.

The orientation model similarly wrapped the angular difference but onto 0° - 90°, such that opposite directions sharing the same orientation have no difference. The orientation model is partially inspired by the fact that direction-selective neurons are often mixed with orientation-selective neurons in the early visual cortex (Geisler et al., 2001; Marshel et al., 2011; Ayzenshtat et al., 2016).

#### 2.3.5 Linear Mixed Effect (LME)

We performed LME analysis using the ‘lme4’ package for R studio (Bates et al., 2015) on both human and mice data. All LME analyses were performed with maximum likelihood (ML) rather than restricted maximum likelihood (REML). This is because we sought to compare the relative performance of models with different fixed-effect structures (Meteyard and Davies, 2020).

For human data, we first logit-transformed the dissimilarity matrix by the following formula, to ensure the data followed a normal distribution. Here, 7 is the fixed maxima of the human dissimilarity scale and ε is a small constant that prevents boundary values (0 and 7) from producing infinities under the logit. We transformed according to this formula, where ε = 0.001, x is the original dissimilarity value and z is the transformed value:

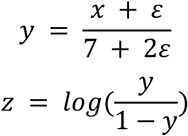

For our LME analyses, we tested the same 3 different direction models in combination with additional fixed-effects. Briefly again, these models were Direction-, Orientation- and Category-based and represent different ways of quantifying the angular difference between stimuli pairs. The Direction model captured absolute angular difference between directions, the Orientation model additionally corrected for symmetry by collapsing opposite directions onto the same orientation axis, and the Category model grouped angular differences into the same four classes: Same, Opposite, Orthogonal and Other. In the case of LME, these models do not change the dependent variable; they differ only in how the same angular relations are coded as fixed effects predicting dissimilarity.

Additional fixed-effect for human data was the absolute value of the difference between logarithm (base 2) of spatial frequency for the stimulus pair (represented on figures as SF). SF was then represented as a factor for the purpose of LME analysis in order to preserve degrees of freedom. Interaction of this fixed effect of spatial frequency with the respective direction models and the random effect of individual participants (1| participant) was also modelled.

For mice data, we identified a single global maximum distance value (MaxVal) across all distance matrices derived for each mouse (in both the awake / anaesthesia condition). This single denominator was used to normalise all distance values, ensuring that mean-level differences between awake and anaesthetised states are preserved since both conditions are divided by the same constant. Then, we transformed all entries of the matrix by the following formula, where ε = 0.001, x is the original distance value and z is the transformed value:

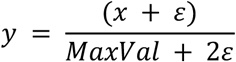

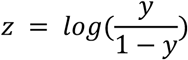

The diagonal entries of the distance matrices for mice data are always 0 due to our definition explained in Section 2.3.2. Accordingly, we excluded these values from all our analyses. For the LME analyses, as in the case of humans, we used the same direction models (Direction, Orientation and Category) and the absolute value of the difference of log spatial frequency. We also considered interactions between direction models, the SFs, wakefulness conditions, and the random effect of individual mice (1| mouseID).

To compare many models in our LME analyses, we conducted a series of nested likelihood ratio tests (Bates et al., 2015), obtaining test statistics (χ^2^) as twice the difference in log-likelihood between the full and a reduced model lacking the fixed-effect of interest. Degrees of freedom (df) are equal to the difference in the number of parameters between the two models.

#### 2.3.6 Polynomial fits to angular difference profiles

For each of the three control experiments (§2.1.7), mean dissimilarity as a function of angular difference was fitted with two nested second-order polynomials via ordinary least squares (lm in R). The unconstrained model (y ∼ x + x²) allowed the parabola to peak at any angle. The symmetric model (y ∼ (x − 90)²) constrained the parabola to be symmetric at 90° and equal at 0° and 180°, which is the prediction of an orientation-only tuning account. The two models were compared by F-test on the residual sum of squares; a non-significant F indicates that orientation-only tuning is statistically sufficient to describe the angular-difference profile, while a significant F indicates residual direction-specific structure beyond what orientation can capture. The vertex of the unconstrained parabola (−β₁ / 2β₂) was used as a descriptive index of where dissimilarity peaks; orientation-only predicts 90°.

## 3. Results

In this results section, we first describe the raw human dissimilarity matrix and mouse neural distance matrix to highlight salient regularities that appear across the data (Fig. 4). We then use MDS as a descriptive tool to visualise dominant structures in these very high dimension matrices. Because these descriptive summaries can emphasise different axes across data sources, we next collapse the matrices across stimulus parameters to isolated shared components of structure (Fig. 5). Additionally, we report on human control studies to address any potential sampling or stimulus property concerns (Figs. 6, 7). Finally, we quantify these observations using linear mixed effects model comparison across candidate stimulus models (Fig. 8).

**Fig. 4.**
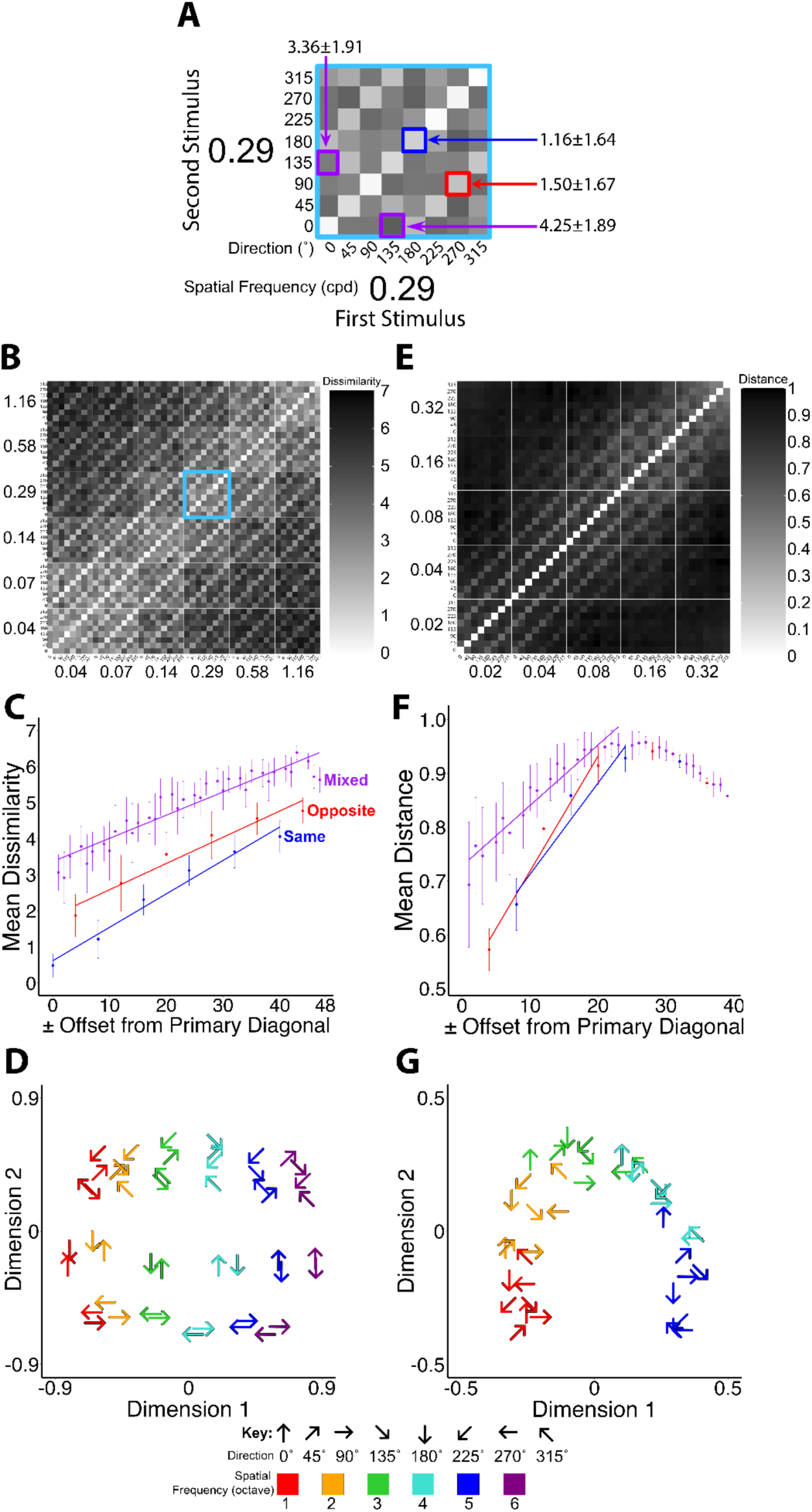
Dissimilarity matrix analysis in humans and distance matrix analysis in mice. Side by side analysis of human dissimilarity data (**A-D**) and mouse neural population activity distance (**E-G**). In all matrices, row and column order is organised by spatial frequency and direction blocks in order to make diagonal offsets interpretable. MDS follows the key shown at the bottom, where arrows denote the actual stimulus direction, and colour marks the ascending spatial frequency octaves shown on the matrix axes **A.** A submatrix (spatial frequency = 0.29 cpd) of the entire dissimilarity matrix highlighted by blue square in **B**. Each cell represents the average pairwise dissimilarity for stimuli represented by rows and columns. **C.** Scatter plot corresponding to B of mean dissimilarity along each diagonal pair offset from the main diagonal. Error bars ±1 SD across mean dissimilarity values along the diagonal at the offset of a particular point. Three groups are identified where direction pairs do not differ (blue), are in opposite directions (red) or differ by 45°, 90° or 135° (Mixed, purple). Lines are linear regression of each group. **D.** Two-dimension solution using Kruskal’s non-metric MDS on the symmetrised dissimilarity matrix in B. **E.** Distance matrix of all awake mice. Rows and columns are organised as per B. See methods Section 2.3.2 for a detailed description of how distance matrices were computed. **F.** Scatter plot of diagonal mean distance corresponding to D as per C. Linear regression lines were computed only for 0.02, 0.04 and 0.08cpd groups (offset = ±24) as mean distance begins to decrease beyond these groups. **G**. Two-dimension solution using classic Euclidean metric MDS on the distance matrix shown in E.

**Fig. 5.**
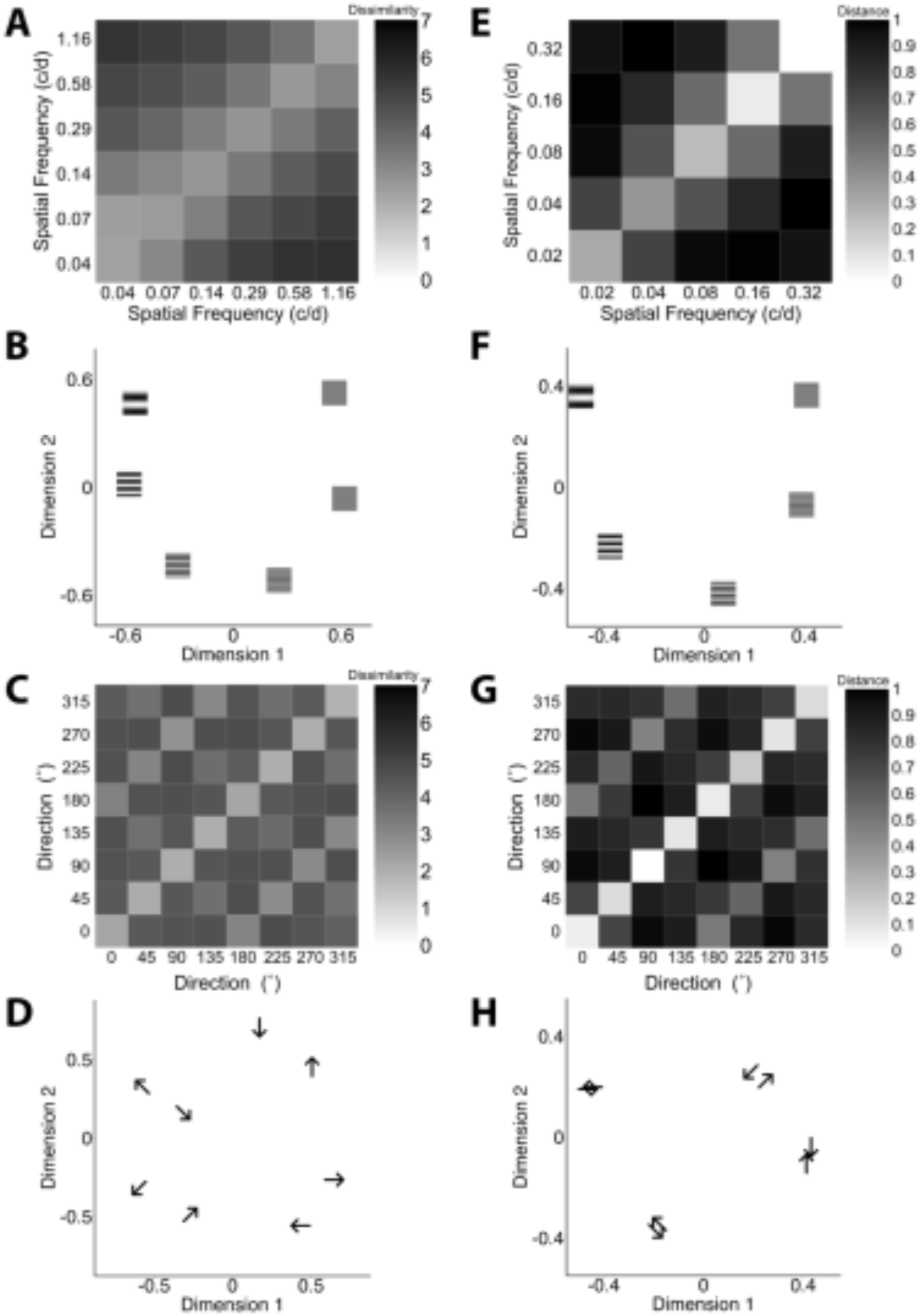
Dissimilarity matrices and MDS collapsed across directions or spatial frequencies for humans and mice. (A-D, Left column) All human data. MDS uses non-metric Kruskal solution in 2 dimensions. (**E-H, Right column)** All mice data. MDS uses classic Euclidean metric solution in 2 dimensions. **A** and **E.** Matrices averaged across directions (in Fig. 4B and Fig. 4E) Each cell represents a spatial frequency group in comparison to another group. **B** and **F.** MDS solution for A and E, respectively, where spatial frequencies are shown as icons on the plot of the spatial frequencies themselves. **C** and **G.** Matrices averaged across spatial frequencies. Each cell represents a direction group. **D** and **H.** MDS solution for C and G, respectively.

**Fig. 6.**
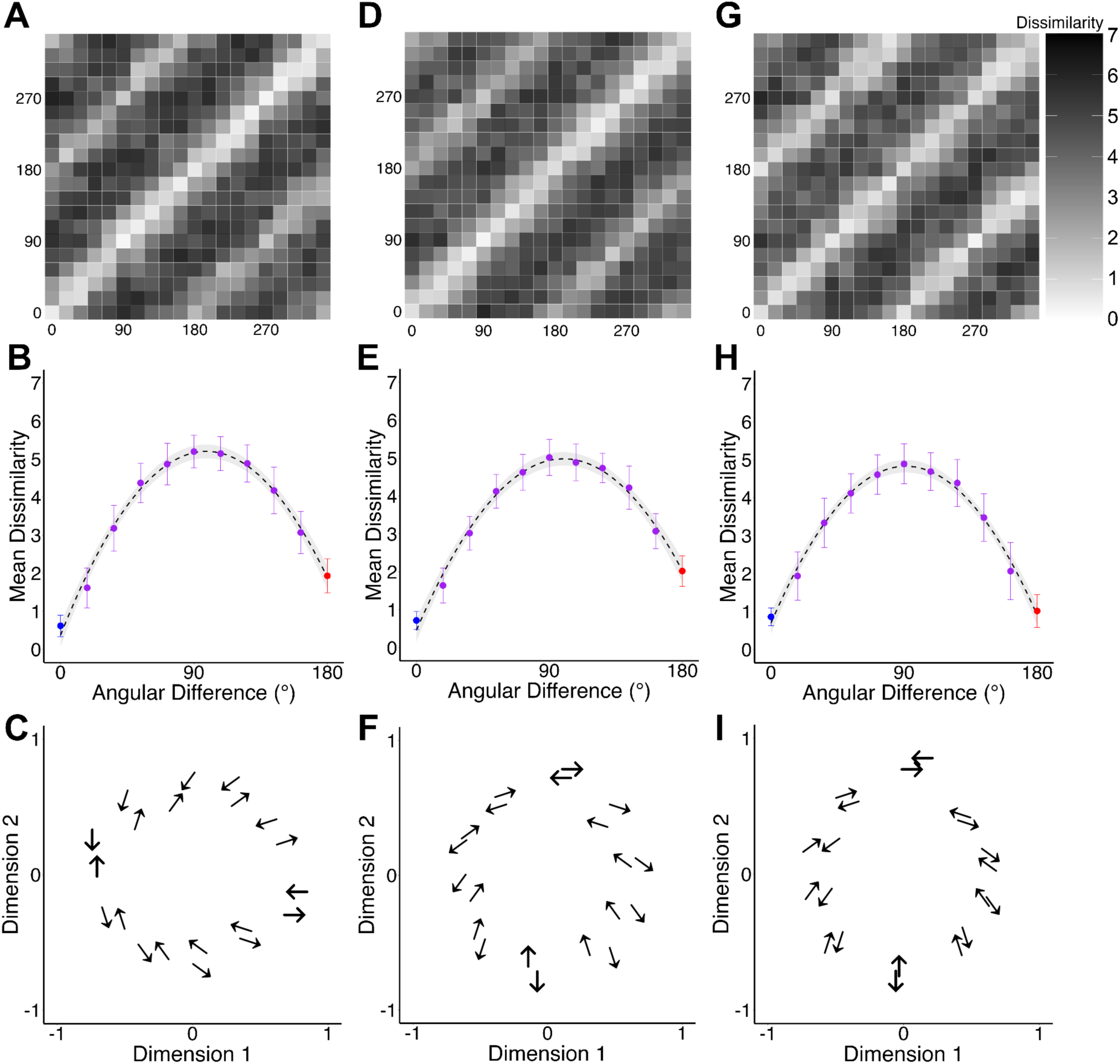
Dissimilarity matrices and MDS from control experiments in humans. Results for visual motion at 0.3Hz (N=16, **A-C**), 2.4 Hz (N=17, **D-F)**, and 18 Hz (N=17, **G-I)**. All dissimilarity matrices (**A, D, G**) are symmetrised within each trial group that had a double-pass correlation greater than 0.139. All matrices use the same colour scale presented next to **G**. Angular difference plots (**B, E, H**) are calculated by averaging along pairs of diagonals that represent increasing angular difference in motion directions. The main diagonal (blue points) represents direction pairs that move in the same direction (angular difference of 0°). Red points represent opposite direction pairs (180°). Purple points represent all other intermediate/mixed degrees of angular difference. Error bars are 1SD across all mean dissimilarity values for that angular difference. Black dashed curves overlaid on B, E, H are unconstrained second-order polynomial fits to mean dissimilarity as a function of angular difference, with 95% confidence bands (grey shading). An orientation-only tuning account predicts a symmetric parabola at 90° with equal values at 0° and 180°; this prediction is statistically consistent with the 18Hz data (H) but not 0.3 and 2.4Hz (B, E). MDS plots (**C, F, I**) all use Kruskal non-metric MDS in two dimensions, where arrows represent the actual direction of the stimuli presented.

### 3.1 Dissimilarity and Distance matrix analysis

To aid in interpretation of the dissimilarity matrices used throughout the remaining results, Figure 3 shows two idealised model matrices composed of the same stimulus parameters as the psychophysics data. Each cell in a dissimilarity matrix represents the mean perceived dissimilarity between a pair of stimuli, with lighter values indicating greater perceptual similarity and conversely for darker values (Fig. 3A&B). The main diagonal is barely dissimilar because each stimulus is compared with itself. Each panel (A/D, B/E) illustrates the pattern of dissimilarity expected if it were driven primarily by continuous direction difference or continuous orientation difference, respectively.

In the direction case, dissimilarity increases continuously with angular separation between motion directions, so nearby directions appear more similar and distant directions more dissimilar (Fig. 3A). In the diagonal offset plot, same direction pairs (blue) are maximally different to opposite direction pairs (red), where mixed conditions are spread evenly between (purple) (Fig. 3D). In the orientation case, dissimilarity depends on grating orientation, such that opposite directions have the same orientation and are therefore indistinguishable (Fig. 3B, E).

#### 3.1.1 Dissimilarity in Humans

Fig.4A magnifies a submatrix at a spatial frequency of 0.29cpd. From this, we make three observations. First, when two identical stimuli are presented one after the other, participants do not necessarily experience them as the same (dissimilarity = 0) (e.g., 180°/180° dissimilarity is 1.16±1.64 for mean±standard deviation (SD) across participants, blue arrow Fig.4A). Second, similarity ratings are not necessarily symmetric (purple arrows, first stimulus at 0° then second stimulus at 135° dissimilarity is 3.36±1.91, while first stimulus at 135° then second stimulus at 0° dissimilarity is 4.25±1.89) (p<0.05, with a paired t-test, t-score = −2.20, df=27). Note that a given pair was tested twice per subject, thus this order effect is due to the within-subject effect. These two observations are common (yet rarely explicitly reported) in similarity experiments using sequential stimulus presentations (Tversky, 1977; Epping et al., 2023). Currently, the underlying psychological and neural mechanisms of these effects in perceptual similarity judgements are not known (See Discussion). Because the order-effect introduces asymmetry in the raw matrix, all subsequent analysis using MDS is performed on symmetrised matrices so that they reflect the underlying pairwise geometry rather than presentation order (see sect. 3.2). We will investigate the asymmetry in a separate future project (for example, using the same approach employed in Epping et al 2023). Third, more specifically to our motion stimuli, opposite direction stimuli are reported as similar. Looking closely at the magnified dissimilarity matrix implies that some opposite-moving gratings were rated very similar to each other (red arrow, first stimulus 270° and second stimulus 90° dissimilarity is 1.50±1.64).

This similarity rating is quite common across almost all oppositely-moving pairs (Fig. 4B). In this full matrix, same direction pairs are on the main diagonals as well as on the sub-diagonals for pairs of different spatial frequencies. For example, a pair for 0° at SF=0.04 cpd is located 8 cells below (or to the left) of the pair for 0° at SF=0.04 and 0° at SF=0.07. Every 8 cells from the main diagonal cells with near 0 (white) dissimilarity, there are sub-diagonal cells with lower dissimilarity from the surroundings, every 8 cells in any direction to the above/below/left/right.

Opposite direction pairs are located 4 cells to the up/down/left/right of the same-direction diagonals. These opposite direction pairs are not as similar to the main diagonals but overall, much more white (more similar) than the surrounding non-opposite direction pair diagonals.

To quantify this impression, in Fig.4C, we grouped all the cells with the same “offset” from the main diagonal together and computed their mean and standard deviations (across pairs, error bars). The same direction diagonals with potentially different spatial frequencies are, thus, located a multiple of 8 offsets from the main diagonals, represented as blue dots in Fig.4C. As expected, these same direction pairs tend to be rated as more similar, but they are clearly less similar when the pairs differ in spatial frequency (the more the offsets).

In this format, it is very clear that the opposite direction pairs diagonals (red), which are 4 + a multiple of 8 offsets from the main diagonal, were rated less similar than the same direction pairs and more similar than the other direction pairs, with comparable spatial frequency differences. The other direction pairs consist of 45°, 90°, and 135° differences in motion direction, with no apparent structure. Thus, we grouped them as “mixed” pairs in purple in Fig.4C.

To statistically verify this visual impression, we performed an ANOVA with Tukey’s HSD on these three conditions (Same, Opposite, Mixed). We confirmed that the dissimilarities of both the same and the opposite diagonals were different from that of mixed diagonals (p = 0.0003 and p = 0.012, respectively), yet those of the same and the opposite were not different (p = 0.44).

#### 3.1.2 Distance in Mice

Next, we looked at neural response patterns obtained from mouse visual cortex activity with equivalent visual motion stimuli. We wanted to see if we would observe a corresponding pattern in mice neural activity to the human psychophysics dissimilarity matrix and its dependency on direction and spatial frequencies. Here, we use the mouse matrix as a representational geometry of V1 population responses; our goal is not to match absolute distance magnitudes across species, but to compare whether stimulus-dependent structure is expressed similarly. Fig.4E and F shows that at lower spatial frequencies (0.02, 0.04 and 0.08 cpd) there is indeed a correspondence. Lighter coloured diagonal lines show relatively lower distance (higher similarity) for opposite direction pairs. For example, at 0.02 cpd, 0° vs. 180° distance is 0.52±0.02 compared to a 0° vs. 90° distance of 0.71±0.02. However, this correspondence breaks down above 0.08cpd. This is likely due to the limits of mouse contrast sensitivity at higher cpd, which decreases around 0.20 cpd, thus the breakdown we observed is consistent with our understanding of mouse visual motion perception (Prusky and Douglas, 2004). Distance matrices constructed for each mouse individually while awake and anaesthetised as well as their difference were also computed (Supplementary Fig. S3).

### 3.2 Structural analysis of visual motion in dissimilarity and neural response patterns

We use MDS here as descriptive visualisations of the dominant organising axes in each matrix. Cross-dataset commonalities are assessed more directly by parameter-collapsed analyses (section 3.2.3) and model-based quantification using LME (section 3.4).

#### 3.2.1 Human visual motion dissimilarity structures

Given the high dimensionality of the dissimilarity matrix of human similarity judgements, we chose to reduce this using multidimensional scaling in order to better visualise relationships in the data. Here, the distance between points in these two-dimensional plots represents their dissimilarity, such that dissimilar points are further away than similar ones.

In humans, MDS in two dimensions (Fig.4D) suggests that the relative experience of directions compared to one another is more or less the same across all spatial frequencies; opposite direction pairs are close together (most similar) for each spatial frequency group. Directions form 3 ‘horizontal’ groups with intermediates 45°, 135°, 225° and 315° in one, 0° and 180° in another and finally 90° and 270°. Additionally, the 6 spatial frequencies form 6 distinguishable ‘vertical’ groups on the plot. Within each SF group, the perception of directions in relation to one another is reasonably well preserved. Each opposite-direction pair is retained in more-or-less the same position in space for each spatial frequency. At a given motion direction, stimuli with a spatial frequency of 0.04cpd are intuitively found most dissimilar at the greatest perceptual distance from those with a spatial frequency of 1.16cpd. This largely reflects the relationship seen in Fig. 4B; compare the spatial frequency row group of 0.04 to each spatial frequency column as they go from light (less dissimilar) to increasingly dark (more dissimilar). The proportion of variance explained by this two-dimension solution is 0.416.

#### 3.2.2 Mouse neural response pattern similarity structures

MDS (Fig. 4G) of the distance matrix representing all awake mice in Fig. 4E is apparently not homologous to the human result (Fig. 4D). While spatial frequency groups do form clear clusters from lowest (0.02cpd) to highest (0.32cpd) moving left to right, within these clusters, there is no clear organisation of directions, resulting in an overall ‘upside down U’ shape. The proportion of variance explained by this two-dimensional solution is 0.44.

The difference between the structures in humans and mice is expected given that the mouse distance matrix is strongly shaped by spatial frequency, whereas human data reveals 3 distinct direction clusters. For this reason, we next examine structure after collapsing across each stimulus parameter to isolate components that can be compared more directly across datasets (section 3.2.3).

#### 3.2.3 Collapsed across stimulus parameters

As MDS solutions presented in Figure 4 revealed differing structures between human and mice data, we investigated these structures further by averaging across each stimulus parameter (direction and spatial frequency). The respective matrices and MDS plots in two dimensions are presented in Fig. 5.

Concerning spatial frequency, the structure of human dissimilarity structure (Fig. 5B) and mouse neural responses (Fig. 5F) are very similar. Both form a ‘U’ shape, from lowest to highest spatial frequency groups running left to right. Individual spatial frequency groups more or less occupy the same relative position between mice and humans in two-dimensional space. This implies that mice V1 neurons respond to pairwise stimuli in a similar pattern as humans subjectively experience those same pairwise stimuli.

Concerning direction, the story is almost the same. In humans, we see that a direction is almost closest (most similar) to its opposite direction pair. This results in 4 groups of those opposite direction pairs (0°/180°, 45°/225°, 90°/270°, 135°/315°). These groups are not as tight as those seen in Fig. 4B, which is expected given the lack of contrast in the dissimilarity matrix (Fig.5C). In mice we also see four groups comprising the four opposite-direction pairs (Fig. 5H). Mice neural responses are still highly similar for opposite-direction pairs and these pairs are much tighter than in humans. The degree to which spatial frequency and direction account for the respective data sets is reported in the section on linear mixed effect analysis.

Overall, when collapsed across spatial frequencies or directions, human dissimilarity structure and mouse neural response structure reveal more common features.

### 3.3 Control studies in Humans

To check whether the opposite-direction similarity effect generalised beyond the main experiment’s design, we conducted three within-subject control experiments using finer direction sampling (18° steps, 20 directions) and a fixed spatial frequency (0.16 cpd), varying only temporal frequency across experiments (0.3 Hz, 2.4 Hz, 18 Hz; see 2.1.7 for full design details and Table 1).

Figure 6 shows the dissimilarity matrices, angular difference plots, and MDS solutions for each experiment. Across all three temporal frequencies, opposite-direction pairs (180° angular difference) were rated as significantly less dissimilar than non-opposite pairs (Wilcoxon signed-rank test, p < 0.05 in all cases), replicating the main finding. Mean dissimilarity for opposite-direction pairs was 1.93 ± 1.40 (0.3 Hz), 2.02 ± 2.10 (2.4 Hz), and 1.01 ± 0.81 (18 Hz).

At 0.3 Hz and 2.4 Hz, opposite-direction pairs were nonetheless significantly more dissimilar than same-direction pairs (Wilcoxon signed-rank, p < 0.05 for both), indicating that participants distinguished opposite from same directions. However, at 18 Hz, same and opposite pairs were not significantly different, and the entire dissimilarity structure became symmetric about 90° — angular differences equidistant from 90° (e.g., 18° and 162°, 36° and 144°) were not significantly different from one another (Wilcoxon signed-rank, all p > 0.05). This was not the case at 2.4 Hz (all such pairs significantly different, p < 0.05) and was mostly not the case at 0.3 Hz (all significantly different except 54°/126°, p = 0.09). MDS solutions (Fig. 6C, F, I) confirmed opposite-direction pairing in a ring-like structure across all three experiments, with the 18 Hz structure showing the strongest 180° symmetry.

To formally test whether the angular-difference profile in each condition is consistent with an orientation-only tuning account (symmetric about 90° and equal at 0° and 180°) we compared an unconstrained second-order polynomial to a polynomial constrained to be symmetric about 90°. At 18 Hz, the unconstrained parabola peaked at 91.2° and the predicted dissimilarities at 0° and 180° differed by only 0.21 units; releasing the symmetry constraint did not significantly improve fit (F(1, 8) = 1.97, p = 0.20), and the constrained model accounted for 98.9% of the variance (R² = 0.989) compared to 99.1% for the unconstrained model (R² = 0.991). At 0.3 Hz and 2.4 Hz, in contrast, the unconstrained parabola peaked away from 90° (98.4° and 99.6°, respectively) with predicted 0°/180° asymmetries of 1.52 and 1.58 units, and the symmetric constraint significantly degraded fit (0.3 Hz: F(1, 8) = 83.4, p = 1.7 × 10⁻⁵, R² dropping from 0.990 to 0.890; 2.4 Hz: F(1, 8) = 93.4, p = 1.1 × 10⁻⁵, R² dropping from 0.989 to 0.863). Together, these tests indicate that the 18 Hz angular-difference profile is statistically consistent with orientation-only tuning, while the 0.3 and 2.4 Hz profiles contain residual structure that orientation tuning alone cannot account for. For implications of these findings, see Discussion section 4.2.

To test whether the axis-like organisation observed in the main experiment reflects orientation alone or a mixture of orientation and direction information, we compared the dissimilarity structure obtained from static oriented gratings (Fig. 7A) to that obtained from drifting gratings in the main experiment (Fig. 7B). Visual inspection of the two matrices reveals broadly similar overall structure, with same-orientation/direction diagonals and spatial frequency grouping preserved in both. However, the key difference is visible in the opposite-direction/orientation sub-diagonal. In the static matrix, pairs separated by 180° are rated as near-identical (as expected, since they are physically the same stimulus), whereas in the drifting matrix, these pairs are similar but not identical.

**Fig. 7.**
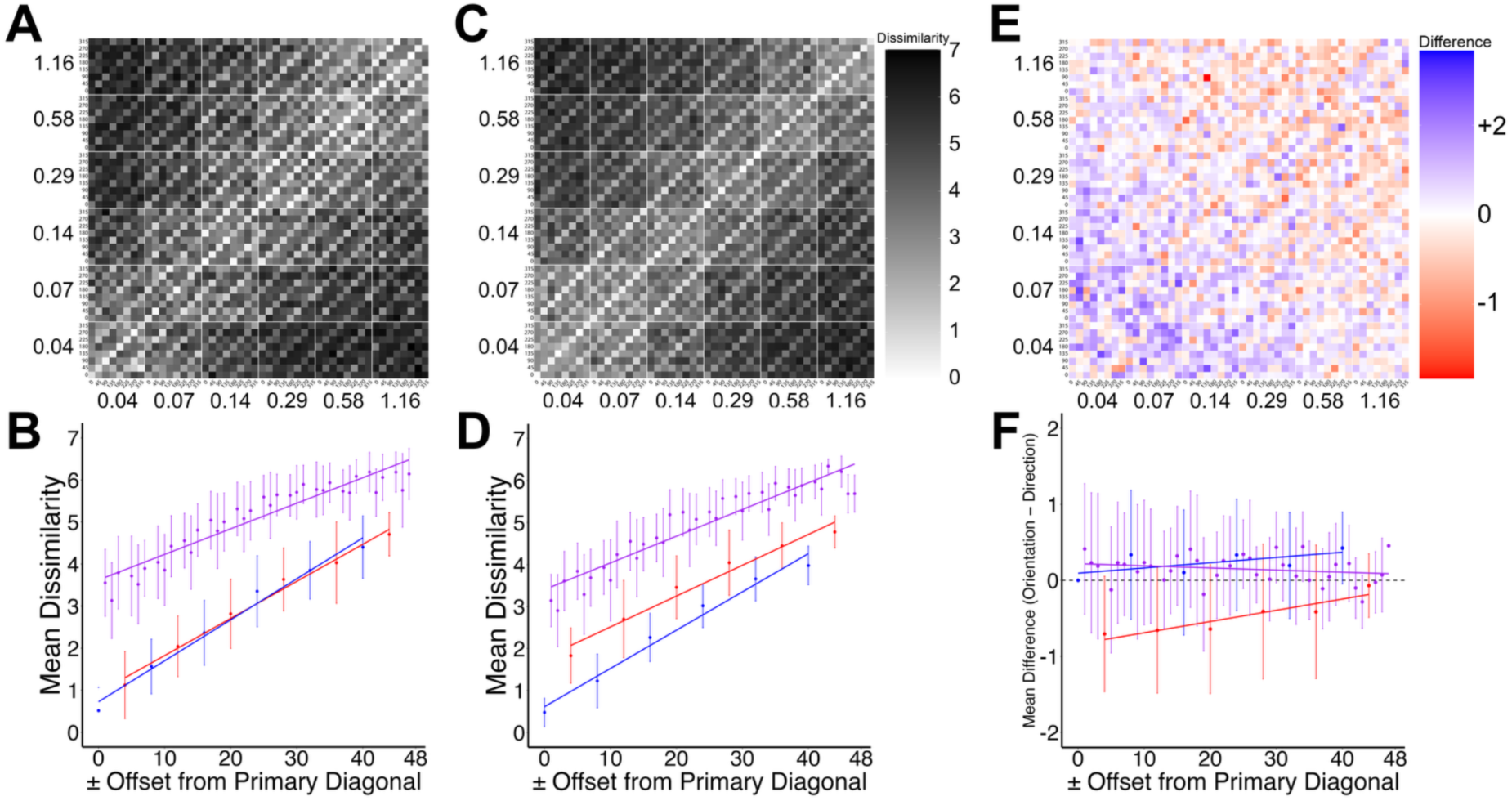
Comparison of orientation-only and direction dissimilarity structures. **(A & B)** Dissimilarity matrix and diagonal offset plot from the orientation-only control experiment, in which participants (N = 81) rated the pairwise similarity of static sinusoidal gratings varying in orientation (8 orientations × 6 spatial frequencies = 48 conditions). Rows and columns are ordered by spatial frequency and then orientation, as in Figure 4B. **(C & D)** Dissimilarity matrix and diagonal offset plot from the main experiment with drifting sinusoidal gratings (N = 171), reproduced from Figure 4 for comparison. Both matrices share the same dissimilarity scale (0–7). **(E & F)** Cell-wise difference matrix (A minus C) and point-wise mean difference in diagonal offset plot. In E, Blue cells indicate stimulus pairs rated as more dissimilar for static than moving gratings; red cells indicate pairs rated as more similar for static gratings. The dominant feature is the red sub-diagonals at opposite-direction offsets (4 + multiples of 8 from the main diagonal), indicating that removing motion collapses opposite-direction pairs to near-identity. All other pairwise relationships show minimal or slightly positive differences (faint blue), indicating that the broader structure is largely preserved. This is represented in F where the difference between the groups of opposite directions in B and D is greater in magnitude than the same or mixed groups which are much closer to 0.

The cell-wise difference matrix (static − moving; Fig. 7E) makes this pattern much clearer. We classified all off-diagonal cells by the directional relationship of the stimulus pair as per the categorical model we used in LME (Same, Opposite, Orthogonal, Other) and tested whether the mean difference in each category deviated from zero.

Opposite-direction pairs showed a mean difference of −0.67 ± 0.90 (one-sample t-test: t(287) = −12.57, p < 2 × 10⁻⁶; Wilcoxon signed-rank: V = 6352.5, p < 2 × 10⁻⁶), indicating that static gratings were rated as significantly more similar (less dissimilar) than moving gratings for these pairs. By contrast, all three non-opposite categories showed small positive differences, indicating static gratings were rated as slightly less similar (more dissimilar): Same (+0.23 ± 0.89; t(239) = 4.01, p = 8.1 × 10⁻⁵), Orthogonal (+0.22 ± 0.84; t(575) = 6.17, p = 1.3 × 10⁻⁹), and Other (+0.19 ± 0.82; t(1151) = 7.93, p = 5.4 × 10⁻⁵). This is more easily visualised in Fig. 7F, which was made by taking the raw mean dissimilarity points at each offset for both orientation and direction matrices and subtracting point-by-point and plotting a regression line from these subtracted points. The mean difference is near zero for Same and Mixed pairs, but substantially greater for Opposite pairs, indicating that motion adds a directional component selectively to the opposite-direction relationship.

A one-way ANOVA confirmed that direction category significantly predicted the difference (F(3, 2252) = 89.05, p < 2 × 10⁻⁶; Kruskal-Wallis: χ²(3) = 211.14, p < 2 × 10⁻⁶). Tukey HSD post-hoc tests showed that the Opposite category differed significantly from all three other categories (all adjusted p < 0.0001), while no pairwise differences were found among Same, Orthogonal, and Other (all adjusted p > 0.91). The overall effect size for Opposite vs all non-opposite pairs was large (Cohen’s d = −1.03; Welch’s t = −15.47, p < 2 × 10⁻⁶). The Opposite-direction effect was consistent regardless of whether the pair shared a spatial frequency (same SF: −0.74 ± 0.81; different SF: −0.65 ± 0.92; t = −0.68, p = 0.50).

These results demonstrate that the drifting-grating similarity structure is not reducible to carrier orientation. Removing motion causes opposite-direction pairs to collapse to near-identity (as expected, since they become the same physical stimulus) while leaving all other pairwise relationships largely unchanged. The fact that moving gratings preserve a residual directional distinction within each orientation axis (opposite pairs are close but not collapsed) is precisely the signature of an axis-like organisation in which direction and orientation information are mixed, consistent with the categorical model that best fits the main experiment data (Fig. 8).

**Fig. 8.**
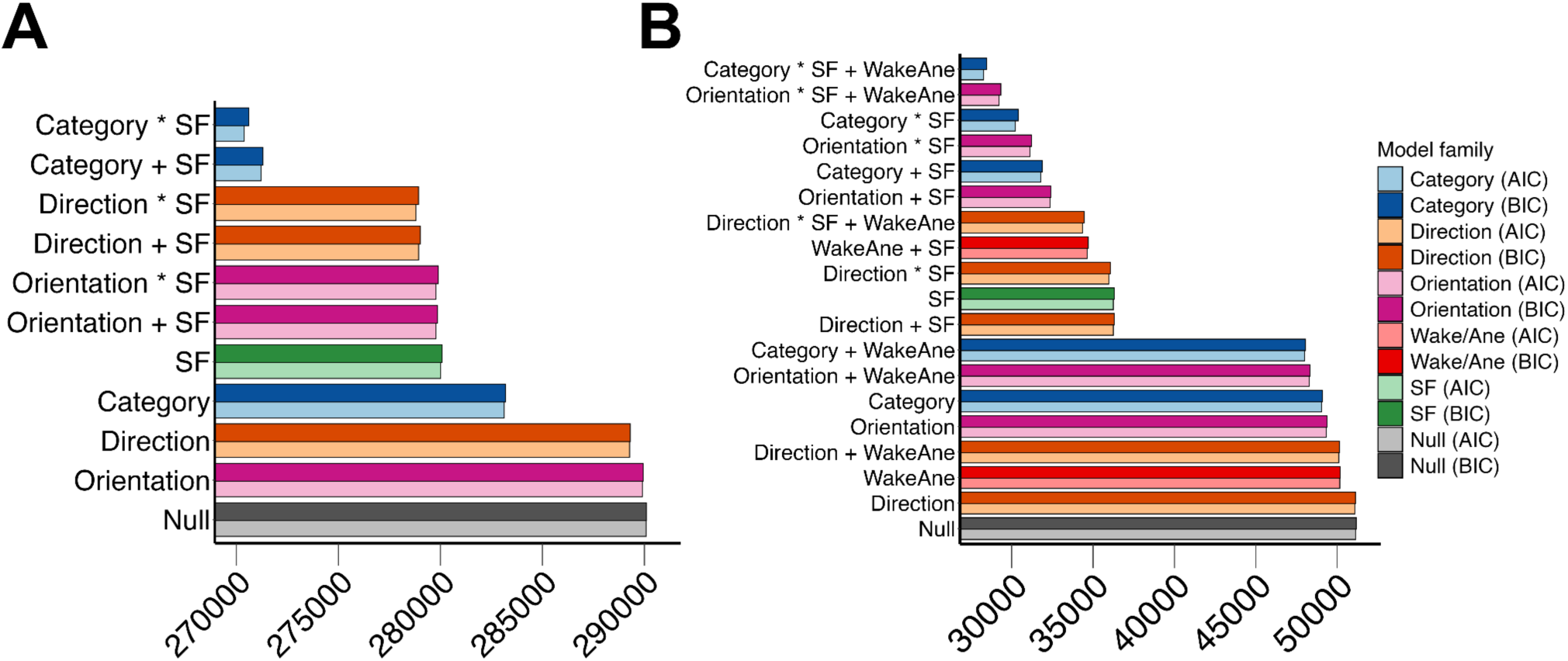
AIC/BIC evaluation of LME models in A) Humans and B) Mice. Relative performance as measured by Akaike Information Criterion (AIC, lighter-coloured bars) and Bayesian Information Criterion (BIC, darker-coloured bars) from LME models in both humans (N=171) and mice (N=9) neuron (n=751) activity. These criterions aim to measure accuracy (AIC) and parsimony (BIC), where lower scores are better. Common to both human and mouse models is the model that has the lowest AIC and BIC is the one that modelled the effects of motion direction as a Category (blue), with a fixed-effect of spatial frequency (SF, green) and its interaction with the motion direction (represented as Category * SF). For the detailed description of three different direction models (Direction (orange), Orientation (pink) and Category (blue)), see Method 2.3.5. Briefly, Category models characterise angular differences in specific categories; ‘Same’ (direction pairs are in the same direction), ‘Opposite’ (direction pairs are in the opposite direction), ‘90’ (direction pairs are 90° apart) and ‘Other’ (all other combinations of direction pairs). Direction and Orientation models predict similarities based on motion directions or grating orientations. **A.** All models contained the random-effect of participants, (1|Participant). **B.** For LME analysis on mice data, all models contained the random effect of individual mice (1|Mouse). Some models contained the fixed-effect of being either awake or lightly anaesthetised (WakeAne, Red) and its interaction with the other factors (motion and SF).

### 3.4 Linear Mixed Effects

Following the qualitative observation that the human dissimilarity structure did not vary monotonically with physical angular difference, and that similar non-monotonic structure was also present in the mouse neural distance matrices, we sought to test these patterns quantitatively using LME analysis. We aimed to test this through three different models (Direction, Orientation, Category) of stimulus direction. A description of these models can be found in Methods section 2.3.5.

Prior to LME analysis, the distribution of participant dissimilarity scores were plotted and logit-transformed as per 2.3.5, then plotted again to ensure the data fit the assumption of normality required for LME analysis (Bates et al., 2015; Meteyard and Davies, 2020).

LME analysis in Figure 8 reveals three common results for dissimilarity based on human judgements and mice neural activity patterns. First, the main effect of spatial frequency (SF) is highly significant (by the AIC, it is −10071.48 and −14903.37 from the null model for human and mice, respectively). Second, the main effect of the Category model is the best among the three models of visual motion (by the AIC, it is −6944.92 and −2094.05 from the null model for human and mice, respectively), compared to Orientation (−172.69 and −1806.87) and Direction (−802.37 and −52.72). Third, interaction between SF and motion models further improves the model fit, thus the full models of Category * SF are always better than other models (−19699.29 and −20915.96). We performed Chi-square tests for each of these model effects, and all were found highly significant (all p values < 10^ij#^ for χ^2^ values greater than 134). A detailed description of the results of these tests can be found in supplementary tables (Table S1, Table S2).

In addition, for mice, the main effect of Wakefulness is significant (Difference in AIC from Null = −981.08) and its full interactions with SF and Category explains the data best (−22868.47).

These statistical results support impressions from qualitative analyses provided in Figures 3-5. With respect to the Categorical motion model (the best fit case), the intercept corresponds to stimulus pairs moving in the same direction and therefore provides the *“same”* as baseline (β = −0.80, 95% CI = [−0.85, −0.74]). Perceived dissimilarity rises by +0.50 (95% CI = [0.47, 0.53]) for opposite motion for humans and decreases by −0.23, 95% CI = [−0.28, −0.19] for mice. It further increases by +0.90 95% CI = [0.88, 0.93] and +0.55, 95% CI = [0.51, 0.59] when directions differ by 90° for humans and mice, respectively. Finally, it further raises by +1.01 (95% CI = [0.99, 1.04]) and +0.35, 95% CI = [0.32, 0.39]) for all other angular separations, for humans and mice, respectively. In sum, these statistical results back up the claim that human perceptions and mice neural activities are 1) most similar when two moving stimuli are in the same direction, 2) the second most similar when they are in the opposite directions, 3) the third most similar when they are orthogonal, and 4) least similar when they are in 45°or 135° difference in directions.

For mice data, wakefulness did affect the neural activity similarity matrix, but its effect was surprisingly smaller compared to the effects of visual properties of the stimuli (SF or motion category). We consider potential implications of this finding in Discussion.

## 4. Discussion

### 4.1 Summary of Results

In this study, we aimed to make a first quantitative connection between a dissimilarity structure used as a method-relative proxy for the structure of visual motion qualia and a mouse neural activity structure. In estimating a human qualia structure from explicit similarity judgements, we sought to provide a more comprehensive understanding of visual motion qualia that goes beyond traditional binary judgments (seen vs. unseen, same vs. different, left vs. right etc). Somewhat counterintuitively, human qualia structure of visual motion presented an axis-like organisation for drifting gratings, in which opposite directions are judged next-most similar to same-direction pairs. Participants tend to experience a pair of opposite directions of motion to be more “similar” than pairs of diagonal or orthogonal motions (Figure 4-6). We found a corresponding neural activity structure in mouse V1 (Figs. 4&5), where opposite motion direction pairs induced more similar patterns of neural responses than other motion direction pairs.

We observed a counterintuitive mismatch between physical angular difference and the inferred qualia structure induced by gratings motion. That is, the most physically distinct opposing direction pairs appear much more similar than physically similar 45°-difference pairs. This is reminiscent of a case of colour qualia structure and light’s physical wavelength structure. A red quale, which can be induced by monochrome wavelengths between 620-750 nm, is experienced as more similar to a violet quale, induced by much more distant wavelengths around 380-420 nm, compared to a very distinct green quale, which is physically closer in wavelength but phenomenally more distinct (Helm, 1964; Zeleznikow-Johnston et al., 2023; Moriguchi et al., 2025). Finally, we found that the V1 activity structure measured in mice was not strongly disrupted by 0.6-0.8% isoflurane in air (Supplementary Fig. S5). We interpret this result carefully; at most it suggests that the coarse V1 response geometry measured here is relatively preserved under this anaesthetic protocol. This finding, in and of itself, is insufficient to infer any conclusion on conscious visual perception.

### 4.2 Opposite directions of motion appear more similar than other directions

As far as we are aware, there is no precedent for attempting to characterise visual motion qualia through similarity ratings. Our findings in both the main (Figs. 4 & 5) and control (Fig. 6) studies show that the inferred qualia structure of visual motion does not directly reflect the physical similarity relations among motion directions. A straightforward alternative explanation is that the qualia structure of visual motion qualia purely coincides with that of visual orientation qualia, largely due to the fact that we used grating as motion stimuli. However, our orientation-only control experiment (Fig. 7) shows a pure orientation account is not sufficient. In fact, removing motion collapses opposite-direction pairs to near-identity while leaving all other relationships unchanged. The observed structure is therefore better described as a mixture of orientation- and direction-related organisation, consistent with the categorical model that best fits both human and mouse data in the LME analysis (Fig. 8). Thus, the most parsimonious description we offer here is therefore an axis-like organisation in which opposite directions share representational structure without becoming identical.

How, then, can such an axis-like organisation be situated relative to classical motion-perception literature? A large body of work in non-human primates has long established area V5/MT as a central component of the motion-perception system through lesion, microstimulation and simultaneous psychometric-neurometric studies (Newsome and Pare, 1988; Newsome et al., 1989; Salzman et al., 1990; Nichols and Newsome, 2002; Born and Bradley, 2005; Clark and Bradley, 2022). Precisely how a neural substrate corresponds to experience is naturally far beyond the scope of this paper, but here we aim to offer a plausible explanation as to how the axis-like dissimilarity structure we observe in visual motion experience could arise from the kinds of neural representations emphasised in this literature. One possible explanation lies in the axis-of-motion columnar architecture of area MT. Albright et al. (1984) showed that macaque MT neurons are organised into continuous columnar slabs within which the preferred axis of motion (the orientation of the motion path) varies smoothly (Albright et al., 1984). Critically, neurons preferring opposite directions of motion occupy adjacent columns within each axis-of-motion slab, so that a full 180° range of motion axes is represented within approximately 400-500μm of cortex. As immediate cortical neighbours, they share much of the circuitry that characterise motion opponency (Qian et al., 1994; Heeger et al., 1999). This columnar arrangement has since been confirmed in human hMT+ using ultra high-field fMRI (Zimmermann et al., 2011; Pizzuti et al., 2023). Additionally, Schneider et al. (2019) showed that columnar clusters in hMT+ track the consciously perceived axis of motion during ambiguous bistable displays (Schneider et al., 2019).

If the structure of explicit similarity judgements reflects, at least in part, the representational geometry of the underlying neural populations (Fink et al., 2021), then this axis-of-motion architecture provides one possible reason why opposite directions should then cluster together in experiential similarity space. Directions sharing a motion axis could activate overlapping cortical territory and opponent interactions within that territory structurally couple opposite-direction representations. From this perspective, motion opponency may not simply sharpen direction estimates (which is often emphasised by classical models of V1-to-MT computation (Simoncelli and Heeger, 1998; Nishida et al., 2018)); it may also bind opposite directions into a shared representational axis. A naïve reading of average-signal motion opponency where the opposite direction is merely subtracted out, would predict maximal dissimilarity between opposite directions. However, at the population level the key suggestion is that opposite-direction neurons could co-localise within axis-of-motion columns, meaning that a readout sensitive to population geometry rather than to the peak firing of individual units would register their structural coupling and thus their relative proximity in representational space. More recently, Chetverikov and Jehee (2023) used a generative model-based fMRI decoding technique to recover full probability distributions of motion direction from trial-by-trial cortical activity in human V1-V4 and hMT+ (Chetverikov and Jehee, 2023). The decoded distributions were bimodal with a primary peak at the true direction and a secondary peak at the opposite direction. Critically, the relative heights of these peaks predicted both the direction and magnitude of behavioural errors, and a parallel bimodality was observed in the distribution of behavioural responses under high uncertainty. These results suggest that the cortical representation of motion direction could link a given direction with its opposite, which is what one would expect from this axis-of-motion architecture. Our own control data are consistent with this account; compared to the lower temporal frequencies (0.3Hz and 2.4Hz), at 18 Hz (corresponding roughly to 112.5°/s) the human dissimilarity structure became more symmetric with respect to 90°, and opposite-direction pairs were rated very similar as if they are same-direction pairs (section 3.3). Under these high-speed conditions, judgments collapsed toward an orientation- and axis-symmetric organisation. A further test would be to ask whether the *degree* of bimodality in decoded cortical representations correlates across individuals with the *degree* of opposite-direction clustering in explicit similarity judgements. If the axis-like structure of motion qualia reflects the geometry of underlying motion representations rather than an idiosyncrasy of the rating task, the two should covary.

Finally, this dissociation between simple angular difference and the estimated qualia structure we present here ultimately needs to be explained within any theory that aims to relate consciousness to physical substrates. One such theory, Integrated Information Theory (IIT) (Albantakis et al., 2023); posits that an experience is identical to the system’s cause-effect structure: the maximally irreducible set of distinctions and relations specified by the substrate in its current state. The axis-of-motion columnar architecture described above offers one possible substrate through which IIT may account for the perceived similarity of opposite directions. Because neurons preferring opposite directions occupy adjacent columns within the same cortical slab (Albright et al., 1984; Pizzuti et al., 2023), they share dense local connectivity that plausibly generate highly overlapping subsets of the full cause-effect structure. It would be interesting to test this prediction by computing model cause-effect structures that incorporate realistic axis-of-motion columnar topography and opponent wiring. Due to various assumptions and flexibilities in possible parameter spaces that need to be tested for such computation (Leung et al., 2021), we believe such a project would be well suited to the Registered Report format (Nosek and Lakens, 2014; Rowe et al., 2024; Cogitate Consortium et al., 2025; Leung et al., 2025), both to constrain analytic flexibility and to ensure informative null results are reported alongside positive findings.

### 4.3 The depth of anaesthesia and the role of murine V1 in visual motion perception

Prior to conducting our analysis, we expected that the human dissimilarity matrix would more closely resemble the neural activity matrix in V1 recorded from wakeful mice than from anaesthetised mice. Yet, we did not find such a clear pattern (Supplementary Fig. S4). The human dissimilarity structure was more or less similar to the structures obtained from neural activity in both states. Spearman correlation between the human dissimilarity matrix (Fig. 4B) and awake mice distance matrix (Fig. 4E) is 0.584, and between human and anaesthetised mice is 0.635. A paired Wilcoxon signed-rank test comparing Spearman correlations to humans between awake and anaesthetised states across individual mice showed no significant difference (p=0.25). Amidst a slew of possibilities, we elaborate two possible explanations on this finding.

First, 0.6–0.8% isoflurane in air may not have abolished consciousness in mice. While this level abolishes behavioural responsiveness and is classically considered sufficient to induce loss of the righting reflex (Nelson et al., 2002; Jurd et al., 2003; Katayama et al., 2007), more recent studies report that LORR requires 1.4–1.5% (Bharioke et al., 2022; Cai et al., 2023), and 1% isoflurane is needed to decouple cortical pyramidal neurons (Suzuki and Larkum, 2020). That said, behavioral unresponsiveness may not necessarily correspond to loss of consciousness (Alkire et al., 2008; Sanders et al., 2012; Zalucki and Van Swinderen, 2016). This could explain why neural activity structures were quite similar between those obtained from awake and anaesthetised (yet potentially conscious) mice. The level of 0.6-0.8% isoflurane in air was chosen as it guaranteed an absence of behavioural response in mice (Wu et al., 2026). A higher dosage, while potentially more effective, would introduce eye drift artefacts that preclude reliable visual stimulation.

Second, V1 representational geometry for these stimuli may be robust to loss of consciousness. In mice, complex visual processing already occurs at the level of V1 (Muir et al., 2015; Palagina et al., 2017; Marques et al., 2018), though perceptual behaviour appears to require secondary visual areas (Jin and Glickfeld, 2020; Goldbach et al., 2021; D’Souza et al., 2022). Consistent with this, Gale et al. (2024) showed that the earliest V1 spikes remain highly informative even when backward masking eliminates perception, suggesting that masking disrupts processing downstream of V1.

### 4.4 Limitations and Future directions

We hope that this study can serve as a proof of concept for this sort of structural analysis in general (Kleiner, 2024), and the qualia structure paradigm in particular (Tsuchiya, 2024). We believe that quantitative comparisons of structures representing phenomenal experience to structures representing a neural substrate will continue to bear fruit and become an increasingly productive and meaningful pursuit in empirical studies of perception. Given this, several limitations and future directions warrant discussion.

A natural first step would be to combine our pairwise similarity-judgement paradigm with human neuroimaging, as recently demonstrated by Hirao and colleagues (Hirao et al., 2025), who constructed a qualia structure for colour perception and compared it with fMRI-derived representational structures. Their use of no-report trials alongside perceptual judgements provides a robust methodology for isolating neural correlates of phenomenal structure without confounds of reporting, and could be applied relatively straightforwardly using our visual motion stimuli. Naturally, remaining within the same species provides direct, first-person access to the phenomenal organisation under study without requiring inference about another organism’s experience. This lends a more useful epistemological constraint on any theory that aims to relate consciousness to a physical substrate, as it must ultimately anchor its phenomenological measurement to the system supporting its experience. Human Neuropixels recordings, now feasible during neurosurgical procedures (Paulk et al., 2022; Leonard et al., 2024), could in principle offer single-neuron resolution, although they require a-priori knowledge of target reasons and are limited by clinical constraints.

Awake behaving non-human primate neurophysiology has been a highly productive avenue for linking neural activity to visual perception for decades (Born and Bradley, 2005), and offers clear advantages for extending the present work. High-density Neuropixels recordings are now possible in macaques (Namima et al., 2024) and marmosets (Dotson et al., 2024), providing well characterised structural and functional homology with human visual cortex, simultaneous behavioural and attentional control, and single-trial population-level analysis during perceptual decisions. These approaches could be used to adequately address the predictions of section 4.2 as large-scale population recordings in macaque MT during motion tasks would be ideally suited to construct neural representational distance matrices and test whether their geometry exhibits the axis-like structure we observe in human similarity judgements. Optogenetic tools for primate cortex (Merlin and Vidyasagar, 2023), including cell-type-specific silencing in MT during direction discrimination (Fetsch et al., 2018), are also maturing rapidly, raising the prospect of causal perturbation experiments in a system with close homology to human motion processing circuitry.

Mouse models offer a complementary set of practical advantages. The extensive genetic toolbox available in mice, including transgenic Cre driver lines for precise cell-type targeting combined with high density electrophysiology across the visual cortex (Siegle et al., 2021), permits a degree of circuit specificity and experimental throughout that remains difficult to achieve in primates, and can be combined with detailed connectomic knowledge of local circuit wiring (Sun et al., 2023). Nakayama and colleagues have demonstrated that mice can perform pairwise similarity judgements with odour stimuli (Nakayama et al., 2022), resulting in dissimilarity matrices that could in principle be coupled with simultaneous large-scale neural recordings (Hong and Lieber, 2019) and optogenetic silencing of target cell populations during the judgement task itself (Gale et al., 2024). However, for both primate and mouse models alike, the fundamental epistemological constraint remains. Relating a neural representational geometry to any structure of phenomenal experience requires inferring subjective experience from behavioural proxies. This gap is certainly narrower for non-human primates, but it is not fully closed for any non-human species, thus a within-species human approach remains the most direct route to anchoring phenomenal measurements.

A further limitation of the present study is the choice of drifting sinusoidal gratings as the visual motion stimulus class. Sinusoidal gratings couple motion direction to orientation, and our orientation-only control experiment (Fig. 7) confirms that the resulting similarity structure reflects a mixture of both, with opposite-direction pairs being the specific locus at which motion adds directional information beyond orientation. Despite this partial resolution, residual effects from the intrinsic coupling remain. As discussed in section 4.2, a critical test of the axis-of-motion account would therefore be to examine whether our results persist with stimuli that decouple direction from an oriented carrier, such as random-dot kinematograms (RDKs) or motion clouds (Moon et al., 2023). In addition, similarity judgements can be systematically shaped by attentional set: recent work shows that feature-based attention can warp perceived feature spaces and their representational geometry (Chapman et al., 2023, 2025), and related dual-task work using similarity judgements suggests that withdrawing attention can qualitatively alter similarity structures for some stimulus classes (Rowe et al., 2025). These considerations together motivate follow-up studies using motion stimuli that better decouple direction from orientated carriers, such as RDKs and/or more naturalistic motion sequences in which objects move against structured backgrounds, consistent with recent proposals that motion perception reflects inference over hierarchical motion relations in richer stimulus environments (Bill et al., 2022).

Finally, two further methodological limitations deserve note. First, techniques for comparing representational structures need to move beyond correlation-based analyses (Kriegeskorte and Kievit, 2013; Roads and Love, 2024), which fail to capture fine-grained structural correspondences. Unsupervised alignment methods such as Gromov-Wasserstein optimal transport (GWOT) (Sasaki et al., 2023), directly address the shape of representational structure without assuming a-priori item-level correspondences or labels. So far, GWOT has been applied to evaluate the degrees of structural correspondence between human groups (colortypical vs. coloratypical) (Kawakita et al., 2023), distinct attentional states in humans (Rowe et al., 2025) and between humans and large language models (LLMs) (Kawakita et al., 2024). Alignment between qualia structures and neural structures, ideally incorporating both anatomical connectivity with activity state (Tsuchiya, 2024), will be a potential new area of research. The coarse correspondence we observed between humans (Fig. 4B) and mouse data (Fig. 4E) suggests that this may work, especially at lower spatial frequencies in mice. Second, our neural activity structures were constructed solely from trial-averaged neural activity, neglecting trial-by-trial fluctuations which may signal causal interactions between neurons. From both a theoretical and empirical perspective, such causal interactions between neurons are strongly suspected as necessary substrates of phenomenal experience (Tsuchiya, 2024; Tononi et al., 2025). While it is possible to target the particular measures of causal interactions based on a theoretical prior (Leung et al., 2021), at this stage, it may be more fruitful to fairly and exhaustively compare all available options of causal interactions, possibly using a Registered Report format (Shimaoka et al., 2024; Leung et al., 2025). Emerging computational toolboxes, such as pySPI, are becoming available to evaluate statistics of pairwise interactions (SPIs) (Cliff et al., 2023; Liu et al., 2025), which includes >250 interaction measures, such as correlation, Granger causality, mutual information, integrated information and many more. Such a project will transparently expose advantages and disadvantages of different neural structures, constructed from various methods (unlike currently dominant correlation based functional connectivity only), and allow us to quantify which types of interactions can possibly align with a qualia structure. Further comparisons between anaesthetised and awake states could reveal which types of neural structures are more or less relevant to levels and/or contents of consciousness.

## 5. Conclusion

To conclude, in this study we present a first step towards comparing qualia structures estimated from human similarity judgements to neural structures estimated from large-scale single neuron recordings from mouse V1. We chose to focus on visual motion qualia and evaluated the similarity of structures between them while mice were awake and under anaesthesia. We found evidence that suggests this line of research is feasible, yet we also identified several issues that need to be addressed in future studies. The qualia structure of visual motion inferred from human similarity judgements indicates that two gratings moving in the opposite directions are rated as more similar than other pairs of directions (excluding identical pairs). Together with our orientation-only control, this pattern is more consistent with an axis-like organisation. This pattern is harder to reconcile with the intuition that perceived similarity should transparently mirror the physical stimulus, and points to the value of characterising experience structurally rather than assuming a direct stimulus–experience correspondence. Structures of neural activity patterns were also unexpectedly similar between those in awake and anaesthetised mice. As such, we regard this study as a proof-of-concept for an across-species structural approach, which holds significant promise, especially with powerful causal techniques (Gale et al., 2024) together with possible similarity judgement experiments in mice (Nakayama et al., 2022). The way that subjective experience can be characterised and quantitatively evaluated as shown here, awaits access to future neural data that has stronger causal links to subjective experience, which is an exciting future indeed.

## Acknowledgements

Thank-you to Davide Aldé and Shelley Smith for their help in editing the final manuscript. NT was supported by National Health Medical Research Council (GNT1183280, GNT2037172), Australian Research Council (DP240102680), Japan Society for the Promotion of Science Grant-in-Aid for Transformative Research Areas (A) (23H04829, 23H04830) and Japan Science and Technology (JST) Moonshot R&D Grant (JPMJMS2295-14), Theoretical Sciences Visiting Program, Okinawa Institute of Science and Technology. AZJ was supported by National Health Medical Research Council (GNT1183280, GNT2037172) and Australian Research Council (DP240102680). YY was supported by Japan Society for the Promotion of Science Grant-in-Aid for Scientific Research (B) (21H026000, 24K02140).

## Footnotes

1 For example, qualia, behavioural reports/responses and/or their underlying neural or functional correlates.

2 Colour perception seems to hold a special place in studies of subjectivity. Consider the age-old question; *is my red your red?* compared to a hypothetical question; *is my perception of something moving up, your perception of something moving up?.* The former seems much more intuitive. Colour seems subjective in a way that visual motion does not. However, it is not fully clear exactly how and in what ways visual colour and visual motion are different from a neural-computation perspective and this leaves the door open to suggest that visual motion perception may not be necessarily veridical or intuitive.

