## Supplementary Material for "Common Phenomenal and Neural Substrate Geometry in Visual Motion Perception"

**Table of Contents**


**Supplementary Notes** **2**
Note on Similarity Structure and Qualia Structure Terminology. 2
**Supplementary Methods** **3**
Distributing Participant Stimuli Assignment 3
Radial response interface for similarity judgements. 3
Tuning properties of the recorded mouse V1 population 4
**Supplementary Results** **5**
Fig. S1. Description of unique stimulus parameters for 171 participants. 5
Fig. S2. Conversion from single mouse V1 neuron's responses to a response pattern dissimilarity matrix. 6
Fig. S3: All mice distance matrices while awake, anaesthetised and their difference. 9
Fig. S4. Spearman Correlation between Human and Mice matrices. 10
Fig. S5: MDS solution for mouse V1 response patterns under anaesthesia 11
Table S1. Likelihood ratio tests of LME models of Human qualia structure. 12
Table S2. Likelihood ratio tests of LME models of Mice V1 neural activity. 13
Supplementary References 14

### Supplementary Notes

**Note on Similarity Structure and Qualia Structure Terminology.**

In this study, the primary human data are pairwise similarity judgments collected with explicit instructions to rate the experienced similarity of two stimuli. From these judgments we construct dissimilarity matrices, which form the empirical basis of all subsequent analyses. In the main text, we use the term qualia structure in the conceptual sense of the Qualia Structure Paradigm: the relational structure inferred from such experienced-similarity judgments. Thus, similarity/dissimilarity structure refers to the directly measured empirical object, whereas qualia structure refers to the inferred experiential structure that these measurements are taken to approximate. We retain the term *qualia structure* in the Introduction and Discussion to align with the broader theoretical framework, while using *similarity* or *dissimilarity structure* in the Methods and Results to describe the data more directly.

### Supplementary Methods

**Distributing Participant Stimuli Assignment**

Out of the full set of 2304 possible stimuli combinations, each participant was allocated a subsample of 150 stimuli pairs for pairwise similarity comparison. We attempted to distribute stimuli to participants in a uniform fashion to ensure equivalent sampling of all stimuli combinations. Specifically, we created an algorithm and database that:

1. Between participants, the current sampling frequency for each stimuli pair in the stimuli comparison matrix was stored in a database.
2. When a participant commenced the experiment, the lowest-sampled stimuli pairs were fetched and allocated to that participant.
3. The stimuli comparison matrix was updated to indicate that those stimuli pairs had their sampling count increased by one, ensuring a new stimuli set would be allocated to the next participant.

Unfortunately, we discovered after the experiment that there was a problem in the algorithm. Specifically, if participants commenced the experiment within a few seconds, before the database could update, participants could be allocated the same set of stimuli. This led to certain stimuli combinations being overrepresented in our sampling matrix (Figure S1).

**Radial response interface for similarity judgements.**

Similarity judgements were collected using an eight-option radial response interface seen in Fig.1C corresponding to integer ratings from -4 to +4 excluding 0. Participants initiated each trial by clicking a small rectangle at the centre of the display and the response interface appeared again after presentation of the second stimulus. In doing so, this configuration requires the respondent's cursor to begin near fixation on each trial before moving to the selected response option.

We have used this interface across many years of large-scale online psychophysics experiments (Qianchen et al., 2022; Kawakita et al., 2023; Zeleznikow-Johnston et al., 2023; Lipeng et al., 2025) and have found it to reduce response bias in online perceptual judgement tasks that use other interfaces such as likert-scales or keyboard button presses. Regarding keyboard presses, we have observed a strong serial response effect, where the button pressed on trial x-1 strongly affects the response on trial x. The radial selection additionally forces that each response option be equal distance (and equal effort) for participants, unlike a likert scale that introduces bias either close to the centre point or to the two extremes (Nicholls et al., 2006).

#### Tuning properties of the recorded mouse V1 population

The 751 neurons (626 excitatory, 125 inhibitory) analysed in this study were recorded via two-photon calcium imaging in layer 2/3 of mouse V1, as described in Wu et al. (2026). All neurons met two inclusion criteria in both awake and anaesthetised conditions: (1) visually evoked responses significantly different from blank (one-way ANOVA, $p < 0.01$), and (2) trial-averaged preferred-stimulus responses exceeding 0.25 $\Delta F/F$. Direction and spatial frequency tuning were characterised by fitting a double-Gaussian (direction) × Gaussian (log SF) model to trial-averaged responses (Wu et al., 2026).

Orientation and direction selectivity in excitatory neurons did not differ between awake and anaesthetised states (OSI: $p = 0.43$; DSI: $p = 0.50$; Wu et al., 2026, Fig. S4A, E), confirming that the features most relevant to the representational distance matrices constructed here remained intact across conditions. Receptive field positions were not individually mapped, as full-field stimuli (~100° azimuth, ~70° elevation) were used in mice to engage all recorded neurons irrespective of RF location.

### Supplementary Results

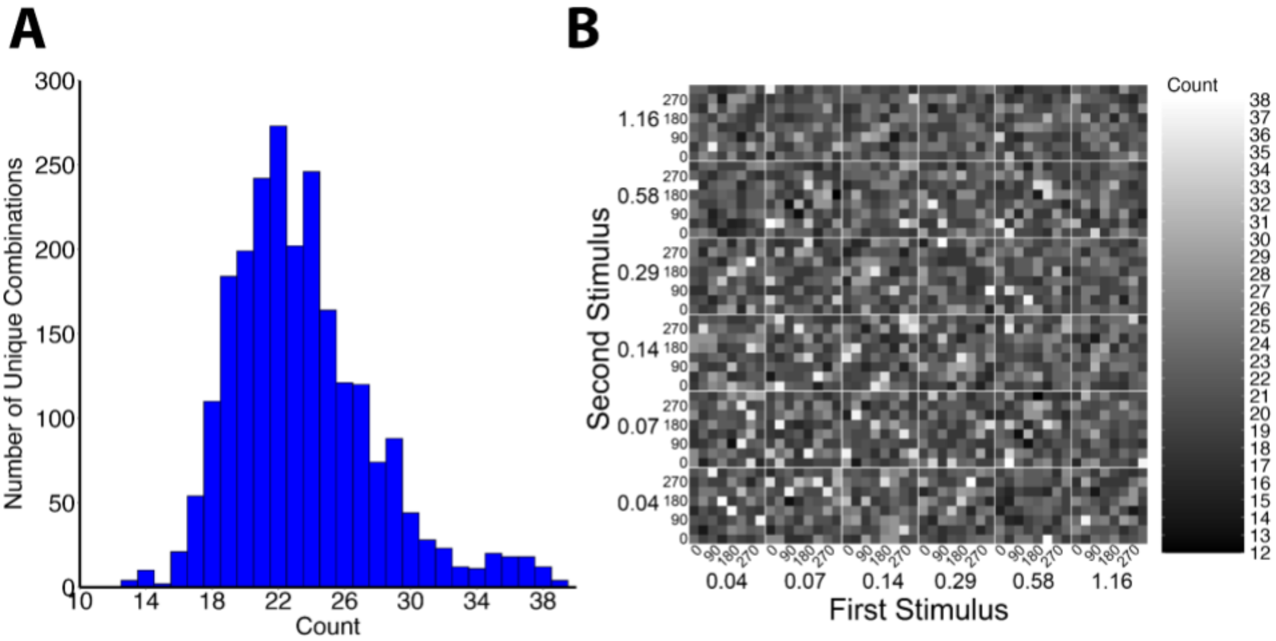


**Fig. S1. Description of unique stimulus parameters for 171 participants.**

There were a total of 2304 unique stimulus combinations which were distributed across participants. **A.** Histogram showing the distribution of these combinations by how often they were presented to participants (count). Count ranges from 12 to 38. Mean count was 22.3 with a variance of 19 and standard deviation of 4.35. Median count was 21. **B.** Matrix representation of the same data shown in A. Here, cells in the matrix represent pairwise stimulus combinations in order of presentation (But for a given subject, we presented the same pair twice between the first and second pass, resulting in the symmetric counts for this matrix). Stimulus combinations represented by white cells were shown in greater numbers to participants and correspond to the right-hand side of the histogram in A.

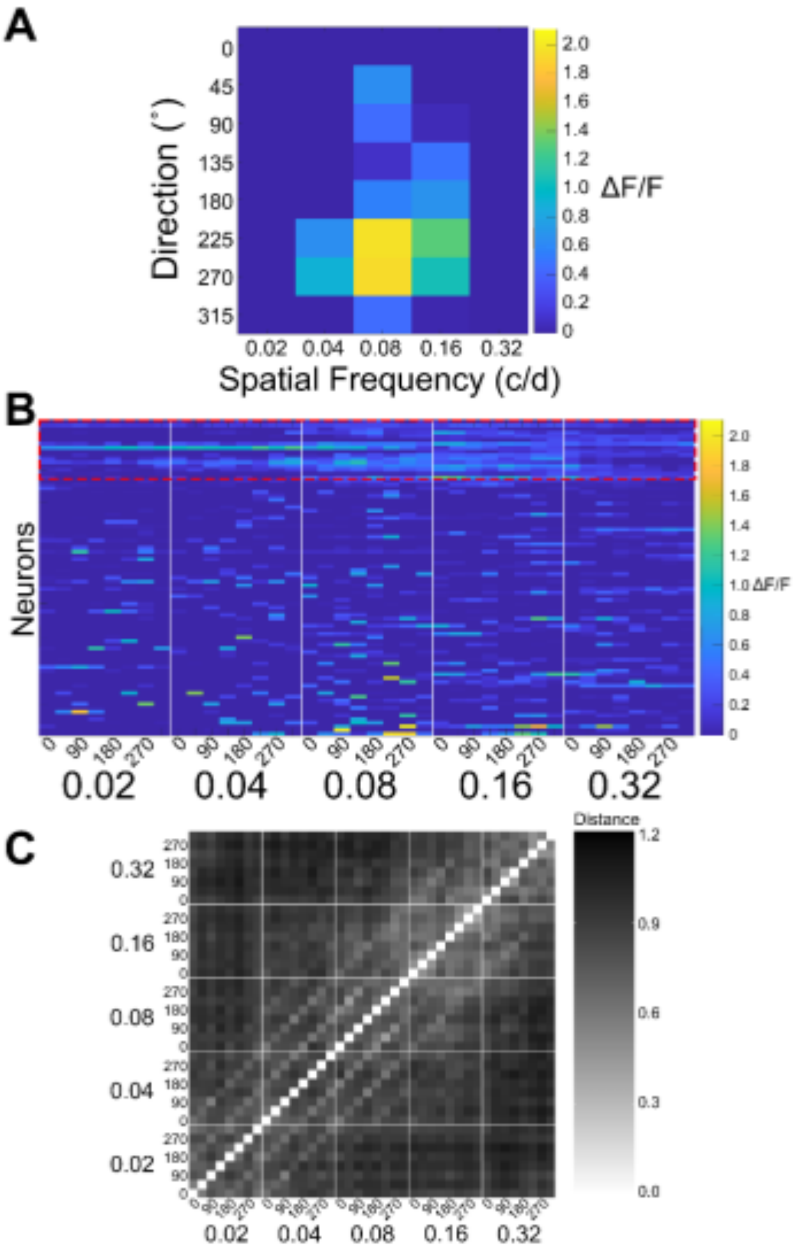


**Fig. S2. Conversion from single mouse V1 neuron's responses to a response pattern dissimilarity matrix.**

**A.** An individual excitatory neuron's activity (neuron ID = 1 in mouse ID = 3) in response to changing spatial frequency and direction of the grating stimuli. Each cell of the matrix represents the neuron's activity in response to stimuli with properties of the combination of its spatial frequency and direction. **B.** All of 85 neurons measured in Mouse 3 (y-axis) as a function of spatial frequency (large numbers) x directions (nested within each spatial frequency, following the format of Figure 4E in the main text). The red dotted box contains all inhibitory neurons (n = 16), the remaining are excitatory (n = 69). The matrix shown in A is flattened to form a single row in B. The neuron shown in A can be found on the bottom row of the matrix. **C.** A distance matrix where each entry of the matrix is $\sqrt{1-r}$, where r is the Spearman correlation value of each column in B to every other column. This process was performed for each mouse while awake and anaesthetised.

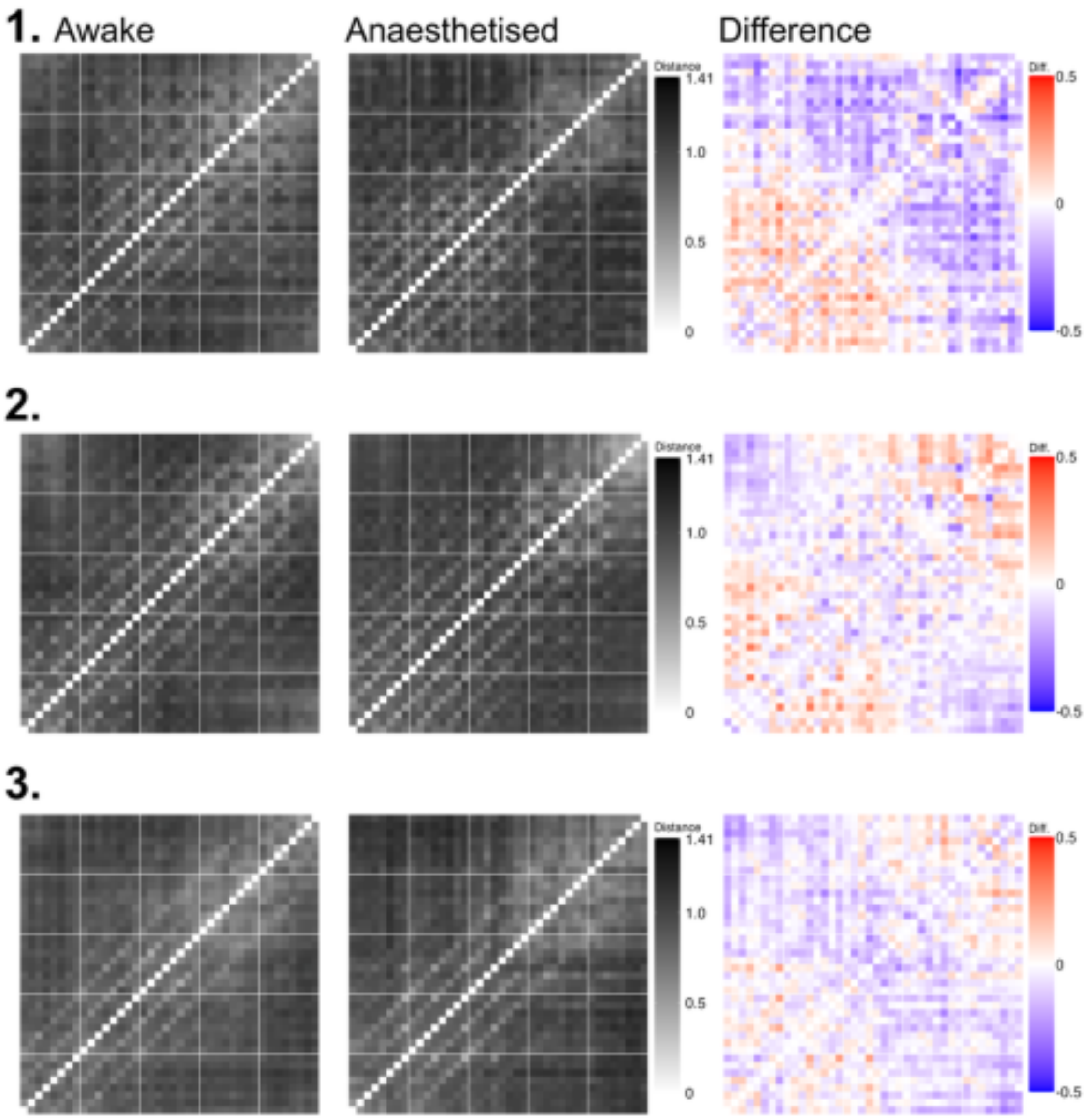
1. Awake
Anaesthetised
Difference
Distance
1.41
1.0
0.5
0
Diff.
0.5
0
-0.5
2.
3.

**4.**

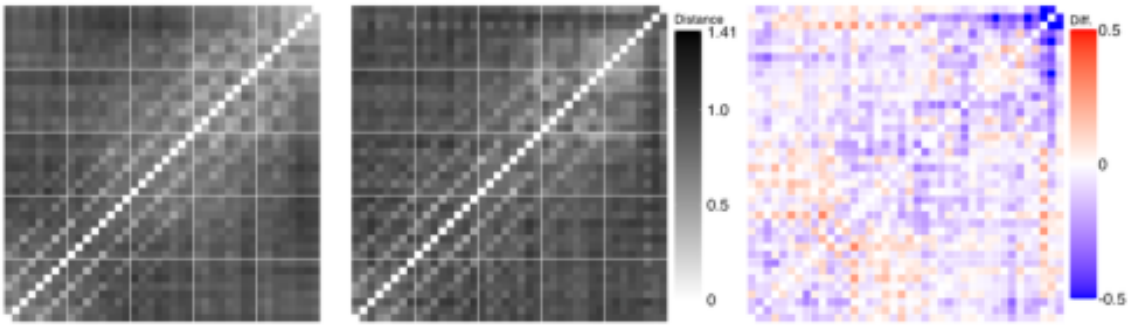
Distance
1.41
1.0
0.5
0
Diff.
0.5
0
-0.5

**5.**

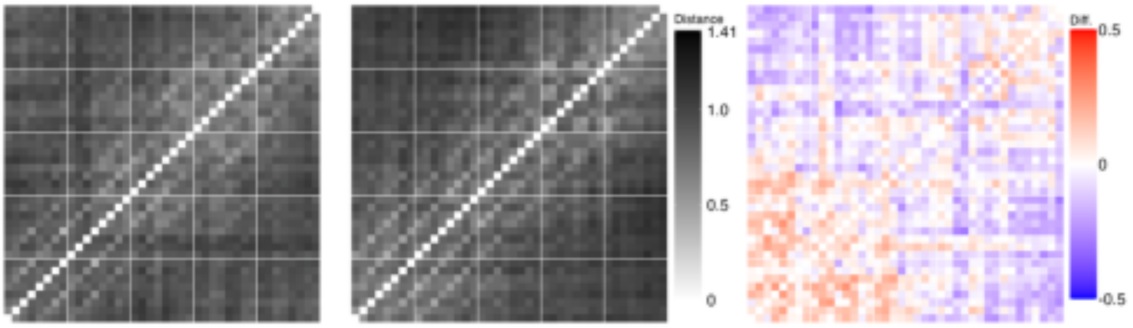
Distance
1.41
1.0
0.5
0
Diff.
0.5
0
-0.5

**6.**

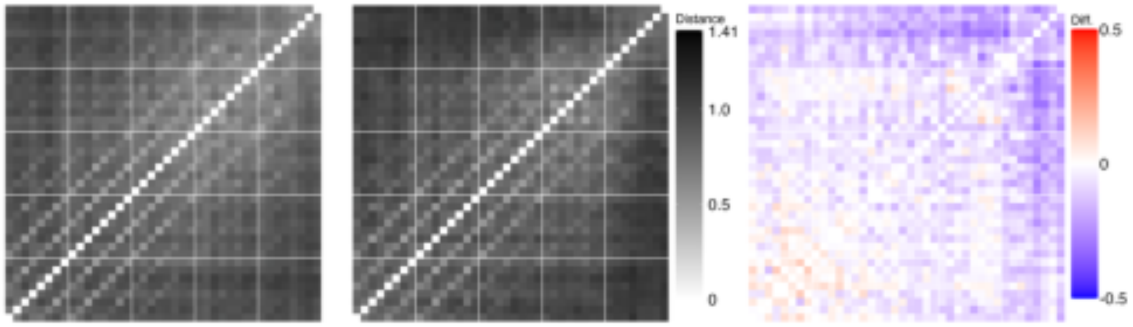
Distance
1.41
1.0
0.5
0
Diff.
0.5
0
-0.5

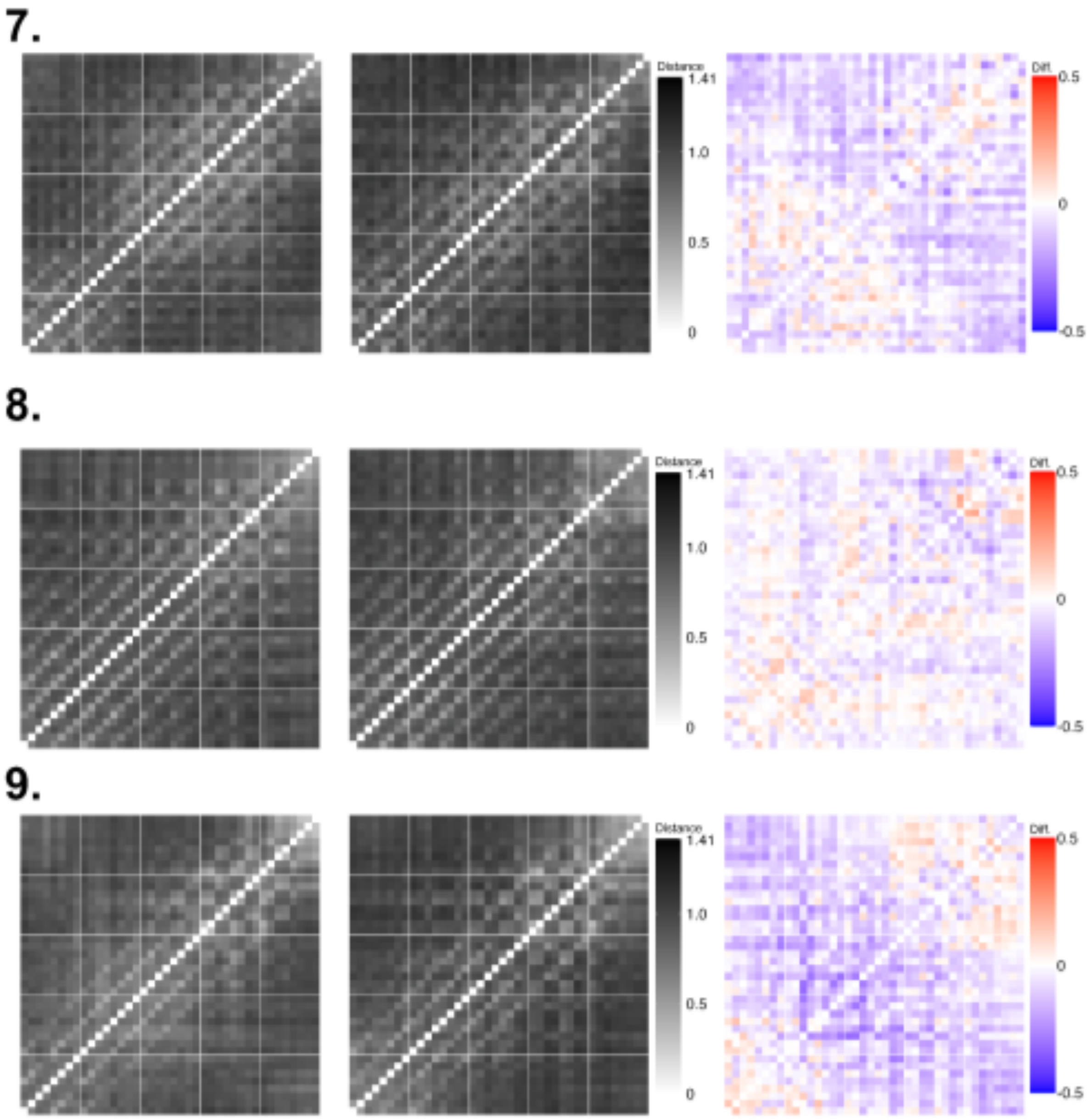


**Fig. S3: All mice distance matrices while awake, anaesthetised and their difference.**

Numbers **1-9** correspond to individual mice ID's. All matrices are composed of data from both excitatory and inhibitory neurons. Axes are unlabeled but follow all previous matrix labels, such as Fig. 4E.

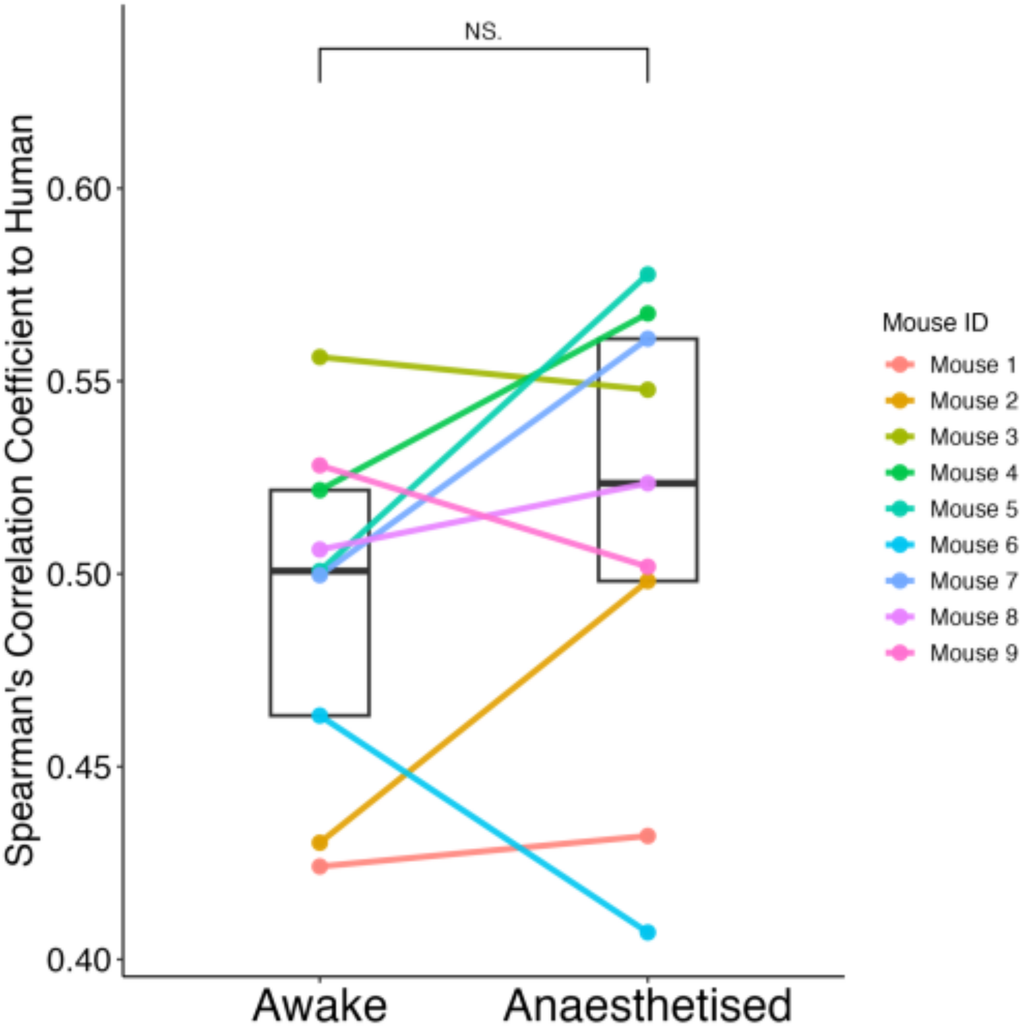


**Fig. S4. Spearman Correlation between Human and Mice matrices.**

Points represent the absolute Spearman correlation coefficient between a single mouse's distance matrix (either Awake or Anaesthetised) and the human reference matrix shown in Fig. 4B. Coloured points and connecting lines indicate individual mice, visualising within-mouse changes between states. Boxplots summarise the distribution of correlations across all mice for each state: the box spans the interquartile range (IQR), the horizontal line marks the median. Significance (NS) indicates the result of a Wilcoxon signed-rank test comparing Awake and Anaesthetised conditions.

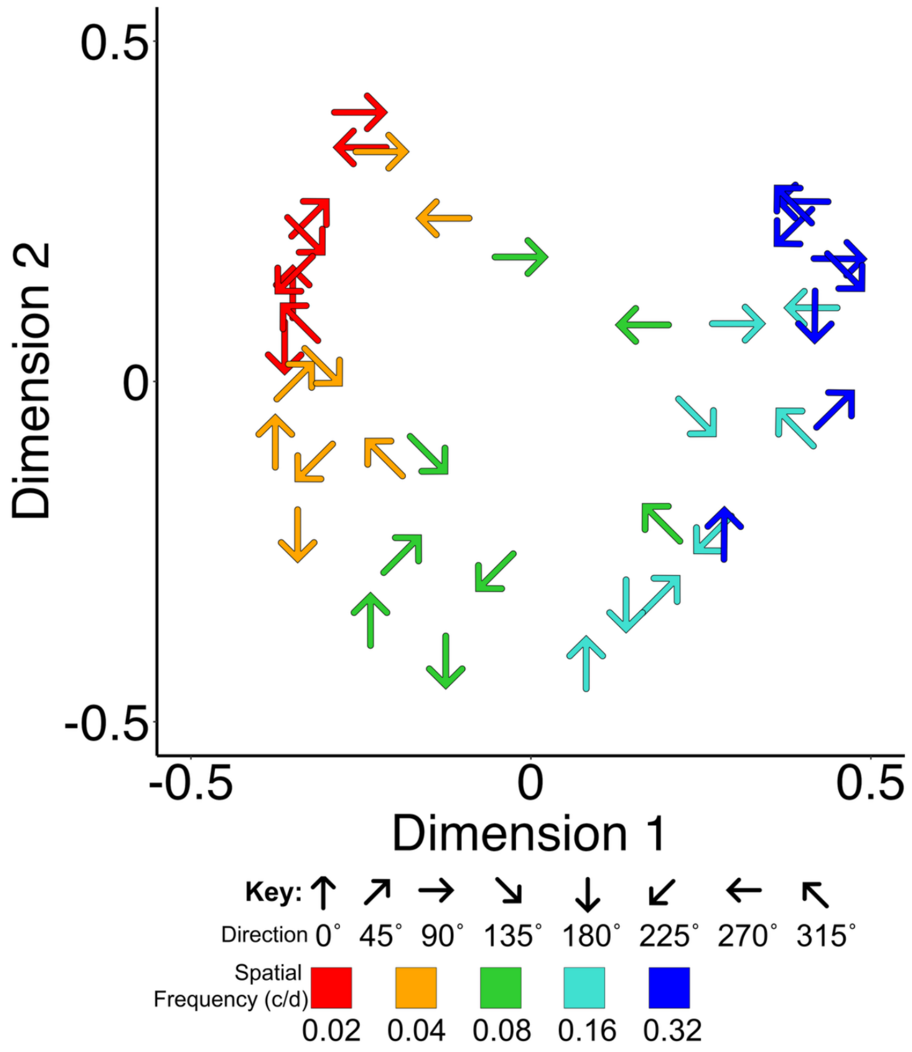


**Fig. S5: MDS solution for mouse V1 response patterns under anaesthesia**

Classic Euclidean metric MDS solution in two dimensions of all mice (N=9) observed neurons (n=751) while anaesthetised.

| **Comparison from:** | **Comparison to:** | $x^2$ | **df** | $p$ |
|---|---|---|---|---|
| (1\| Participant) | (1\| Participant) + SF | 10081.48 | 5 | $< \mathbf{10^{-16}}$ |
| (1\| Participant) | (1\| Participant) + Direction | 804.37 | 1 | $< \mathbf{10^{-16}}$ |
| (1\| Participant) + Direction | (1\| Participant) + Direction + SF | 10351.84 | 5 | $< \mathbf{10^{-16}}$ |
| (1\| Participant) + Direction + SF | (1\| Participant) + Direction*SF | 135.44 | 5 | $< \mathbf{10^{-16}}$ |
| (1\| Participant) | (1\| Participant) + Orientation | 174.78 | 1 | $< \mathbf{10^{-16}}$ |
| (1\| Participant) + Orientation | (1\| Participant) + Orientation + SF | 10130.21 | 5 | $< \mathbf{10^{-16}}$ |
| (1\| Participant) + Orientation + SF | (1\| Participant) + Orientation*SF | 18.50 | 5 | = 0.0024 |
| (1\| Participant) | (1\| Participant) + Category | 6950.92 | 3 | $< \mathbf{10^{-16}}$ |
| (1\| Participant) + Category | (1\| Participant) + Category + SF | 11933.58 | 5 | $< \mathbf{10^{-16}}$ |
| (1\| Participant) + Category + SF | (1\| Participant) + Category*SF | 860.79 | 15 | $< \mathbf{10^{-16}}$ |

**Table S1. Likelihood ratio tests of LME models of Human qualia structure.**

| **Comparison from:** | **Comparison to:** | $x^2$ | **df** | ***p*** |
|---|---|---|---|---|
| (1\| Mouse) | (1\| Mouse) + SF | 14911.38 | 4 | $< \mathbf{10^{-16}}$ |
| (1\| Mouse) | (1\| Mouse) + WakeAne | 983.08 | 1 | $< \mathbf{10^{-16}}$ |
| (1\| Mouse) | (1\| Mouse) + Direction | 54.73 | 1 | $< \mathbf{10^{-12}}$ |
| (1\| Mouse) + Direction | (1\| Mouse) + Direction + SF | 14858.01 | 4 | $< \mathbf{10^{-16}}$ |
| (1\| Mouse) + Direction + SF | (1\| Mouse) + Direction*SF | 278.56 | 4 | $< \mathbf{10^{-16}}$ |
| (1\| Mouse) + Direction*SF | (1\| Mouse) + Direction*SF + WakeAne | 1616.20 | 1 | $< \mathbf{10^{-16}}$ |
| (1\| Mouse) | (1\| Mouse) + Orientation | 1808.87 | 1 | $< \mathbf{10^{-16}}$ |
| (1\| Mouse) + Orientation | (1\| Mouse) + Orientation + SF | 16994.11 | 4 | $< \mathbf{10^{-16}}$ |
| (1\| Mouse) + Orientation + SF | (1\| Mouse) + Orientation*SF | 1233.70 | 4 | $< \mathbf{10^{-16}}$ |
| (1\| Mouse) + Orientation*SF | (1\| Mouse) + Orientation*SF + WakeAne | 1896.24 | 1 | $< \mathbf{10^{-16}}$ |
| (1\| Mouse) | (1\| Mouse) + Category | 2100.05 | 3 | $< \mathbf{10^{-16}}$ |
| (1\| Mouse) + Category | (1\| Mouse) + Category + SF | 17266.09 | 4 | $< \mathbf{10^{-16}}$ |
| (1\| Mouse) + Category + SF | (1\| Mouse) + Category*SF | 1585.82 | 11 | $< \mathbf{10^{-16}}$ |
| (1\| Mouse) + Category*SF | (1\| Mouse) + Category*SF + WakeAne | 1954.52 | 1 | $< \mathbf{10^{-16}}$ |

**Table S2. Likelihood ratio tests of LME models of Mice V1 neural activity.**
